# Leveraging protein representations to explore uncharted fold spaces with generative models

**DOI:** 10.1101/2025.10.10.681606

**Authors:** Yangyang Miao, Martin Pacesa, Sandrine Georgeon, Joseph Schmidt, Tianyu Lu, Po-Ssu Huang, Bruno Correia

## Abstract

A major challenge in computational *de novo* protein design is the exploration of uncharted areas within the protein structural space. However, the large degrees of freedom of protein backbones complicate the sampling process during protein design. Machine learning-based models have made great strides in this problem, however due to their nature they tend to exploit rather than explore the data distribution used for training the neural networks. To address some of these challenges, we propose a new coarse grained protein structure representation generative method, **DiffTopo**, a diffusion model which increases the sampling efficiency and diversity. Combined with a backbone level protein generative model like RFdiffusion, novel protein folds can be generated rapidly, allowing for efficient exploration of the designable topology space. Interestingly, we have discovered that by mirroring the topological organization of native proteins using a pipeline named **MirrorTopo**, we can readily expand the known fold space. We generated and experimentally characterized 30 different novel topologies from DiffTopo and 6 different novel mirror topologies from MirrorTopo. The developed framework relying on low resolution sampling provides new means for fold exploration challenges, which could in principle enhance our knowledge of the first principles of protein structure and folding, as well as create new opportunities for functional design.

## Introduction

Proteins govern biological functions in nature through their three-dimensional structure, yet the vast space of possible folds remains largely unexplored^1^. While natural evolution has populated only a fraction of this space, computational methods now offer a route to systematically chart unexplored structural spaces^2–5^. One of the important challenges in protein design is to efficiently explore the designable protein fold space without being confined to the training data and in this way favor exploitation of that data distribution, rather than exploration. Here, we present a framework that explores the fold space using coarse-grained representations to traverse this landscape, enabling discovery of novel folds. Recent advances in deep generative models, particularly diffusion-based methods, have dramatically expanded the capabilities of de novo protein design. Recently, many deep generative models have been developed that can generate atomically detailed protein structures with complex topologies, such as RFdiffusion^6^, FrameDiff^6^, and Chroma^7^, among many other deep generative models^8–11^. However, despite their success, these methods still tend to sample structures that resemble natural folds, especially for small proteins. This reflects a fundamental limitation of coordinate-level generative modeling: the tendency to recapitulate training distributions and bias the design space toward naturally occurring solutions. As a result, large regions of the theoretically designable but evolutionarily unoccupied fold space, so-called “dark” structural matter, remain underexplored.

We hypothesized that an efficient exploration of the fold space requires a different approach: Rather than sampling structures at the atomic level, the exploration should first occur in a low-resolution topological space where structural diversity and fold novelty is more accessible (Fig. 1a). Drawing inspiration from representations of proteins as hierarchical assemblies of secondary structure elements^12–16^, we introduce a coarse-grained (CG) topology description that describes folds into spatial arrangements of *α*-helices and *β*-strands (Fig. 1b). This reduced-dimensionality manifold smooths the sampling landscape, allowing for more coarse sampling of the structural space and avoiding the combinatorial explosion of atomic representations. We developed DiffTopo, which learns fold topologies through diffusion modeling guided by secondary structure descriptors (a string of secondary structure types), allowing us to sample regions far from natural folds while keeping physical plausibility.

**Figure 1.**
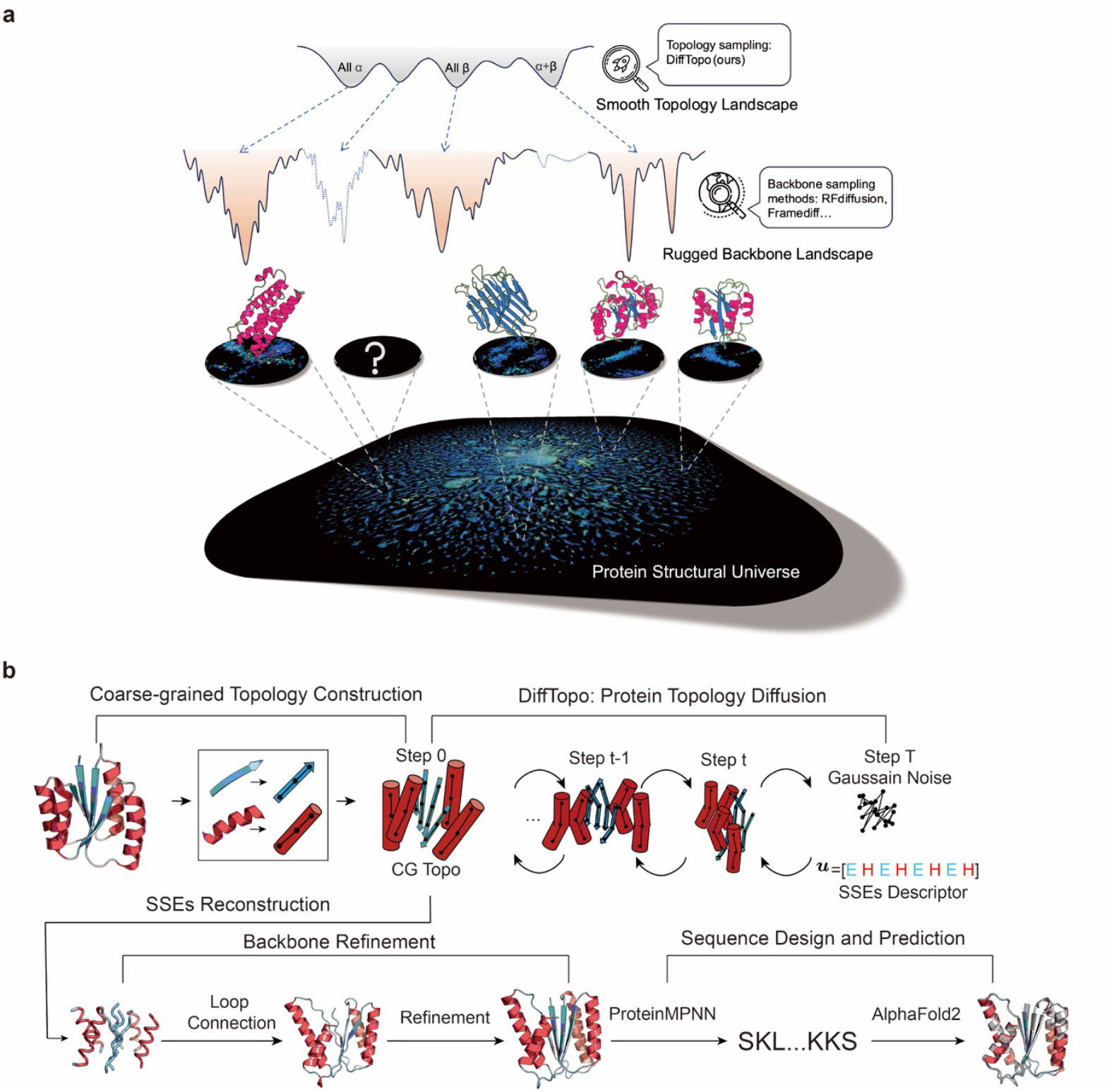
Overview of the protein design landscape and DiffTopo-RFdiffusion pipeline. **a**, Conceptual representation of the Protein Universe Sampling across structural landscapes with different resolutions. The top row illustrates our proposed topology-level sampling approach, which operates at a coarse-grained (CG) level. The rugged backbone-level landscape, where most existing methods, such as RFdiffusion and FrameDiff operate. The bottom row depicts the natural protein universe derived from ESMFold predictions atlas^17^. **b**,Schematic representation of the DiffTopo-RFdiffusion pipeline. The workflow begins with the conversion of secondary structure elements into CG topology representations. DiffTopo is a diffusion model that generates CG topologies that serve as structural blueprints to guide RFdiffusion on the sampling of atomic-level protein backbones.

Coarse-grained topological exploration of protein structure provides an approximate placement of secondary structure elements in 3D space; subsequently, obtaining a full atomic backbone requires a high-resolution generative model.

Based on the CG topology length, we first assign standard secondary structure elements and place them at the corresponding spatial positions, resulting in an intermediate structure composed solely of idealized secondary elements, which we refer to as a *protein sketch*. Our framework bridges this gap by refining these sketches and using them to guide RFdiffusion5 generation, ultimately producing high-resolution, designable protein backbones. Importantly, this separation between fold discovery (DiffTopo), through CG sampling and structure generation (RFdiffusion), via atomic-level modeling, helps overcome some of the limitations of directly generating full atomic coordinates, which often tends to recapture existing topologies from known fold space. We further tested this by attempting a distinct strategy of structural generation, where we systematically generated mirrored image topologies of natural folds, which are absent from natural repertoires but are physically realistic and potentially designable, we dubbed this approach MirrorTopo. These two challenges of generating novel-to-nature dark folds and mirrored topologies demonstrate how coarse-grained sampling can reveal underexplored regions of structural space.

Biochemical and structural validation confirmed the potential of this approach, where 40 designed dark folds and 22 mirror folds were experimentally characterized, with five crystal structures closely matching the computational models. These results suggest that our method can access folds beyond the capabilities of current generative approaches. By decoupling fold exploration from atomic-level sampling, we establish a new strategy to explore protein space that prioritizes discovery over optimization, a critical step toward understanding the full scope of possible protein architectures and their potential functions.

## Results

### Enhancing the sampling of structural diversity with DiffTopo

To test the capabilities of the DiffTopo-RFdiffusion pipeline, we generated 3000 backbone structures for each of the following categories: I) random secondary structure descriptors guided backbones; II) low-frequency secondary structure descriptors guided backbones; III) random lengths ranging from 40 to 200 residues backbone generated by RFdiffusion, as reference. For categories I and II, we describe the secondary structure descriptor guided generation in detail in the methods section. To test whether the CG topology correctly captures the target protein fold and whether the RFdiffusion-generated backbones recover the intended structure, we compared protein sketches derived from CG topology with native structures to ensure the model learns the correct structural features, and with the corresponding RFdiffusion-generated backbones to verify that the generated outputs preserve the intended topology. Most protein sketches match well with the generated backbones (RMSD <3 Å), confirming that the CG sketches can guide the sampling of RF-diffusion for the generation of all-atom protein backbones (Fig. 2a). To assess whether the model effectively learned the geometric distribution of sketch data, we analyzed its geometric descriptors and evaluated their alignment with native structures. DiffTopo-generated sketches match the topological patterns observed in native structures, showing that the model effectively learns an accurate approximation of real data distributions (Supplementary Fig. 1a-c).

**Figure 2.**
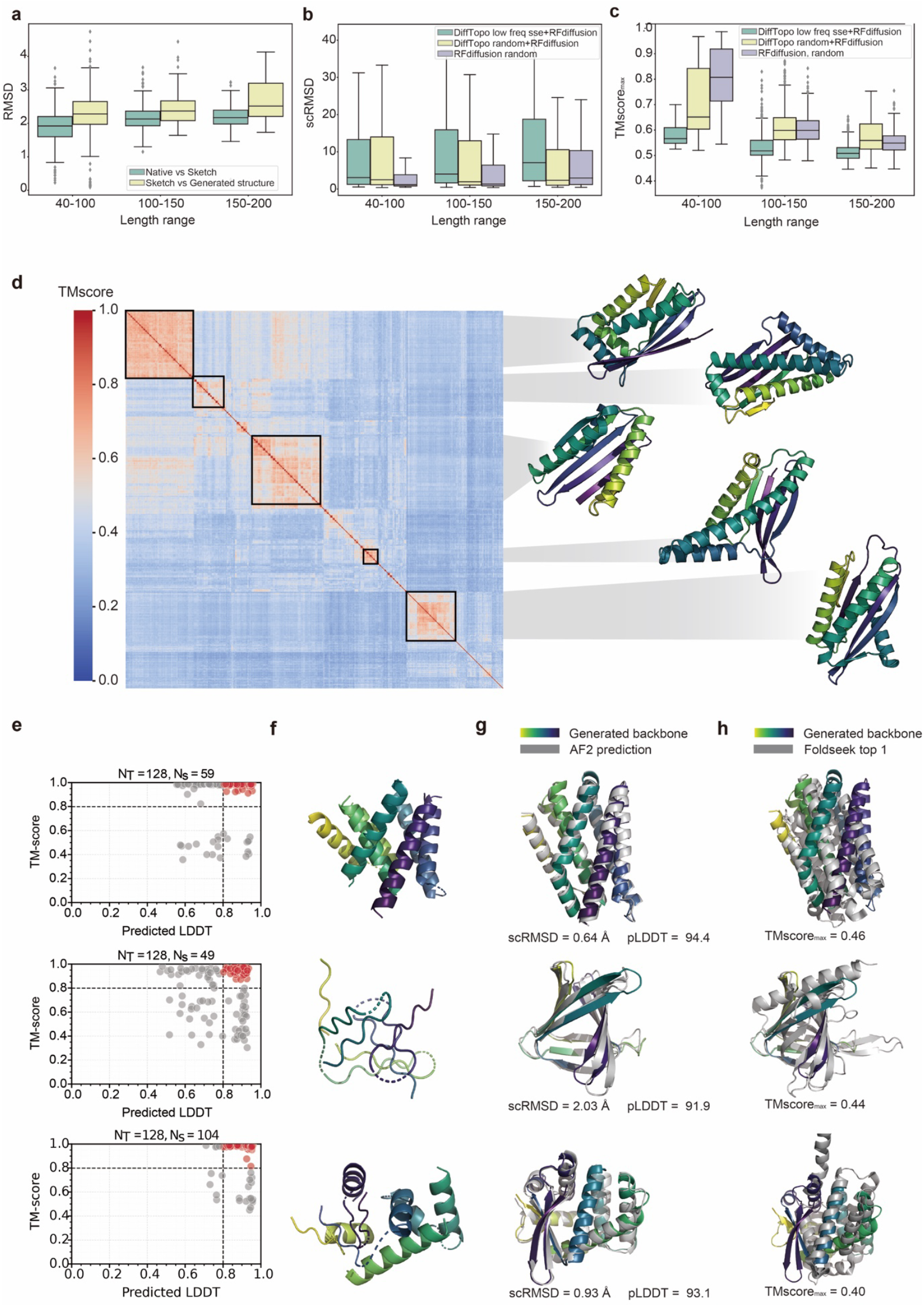
Structural evaluation of DiffTopo-guided sampling. ***a***, Structural comparison between native structures and their corresponding protein sketches from CG topologies (green), and between protein sketches and generated backbones (yellow). Each backbone was generated by RFdiffusion conditioned on the protein sketch. **b**, Designability of generated backbones. scRMSD based on 1,000 samples per length range (40–100, 100–150, 150–200) across different methods. Purple: Rfdiffusion random sampling (baseline), yellow: DiffTopo-RFDiffusion random sampling of SSE strings, green: DiffTopo-RFDiffusion sampling of low-frequency SSE strings in native proteins. **c**, Novelty of generated backbones: color scheme follows b. **d**, Diversity of generated backbones: TM-score heatmap after hierarchical clustering, showing structures from the top 5 largest clusters. **e**, TM-score between DiffTopo-generated backbones and AF2 predictions plotted against predicted pLDDT to assess the designability of the generated structures. Red points indicate highly designable sequences (scTM-score >0.8, pLDDT >80). **f**, Representative visualizations of DiffTopo-generated backbone sketches. **g**, Backbone alignment between DiffTopo-generated structures and AF2 predictions to evaluate the structural consistency. **h**, Comparison of DiffTopo-generated backbones with their nearest structural analogs identified by FoldSeek in PDB and CATH databases. Designs in row 1-3 correspond to N35, N40 and N37 in the Supplementary table 2.

We then assessed the designability of the generated backbones from different types of protocol by designing amino acid sequences using ProteinMPNN^18^ and predicting whether the designed sequences could fold into the target topology with AlphaFold2(AF2)^19^. Following prior studies on protein backbone generation^20,21^, we quantified self-consistency using C*α* RMSD (scRMSD, lower values indicate better consistency) and predicted local distance difference test (pLDDT, higher values indicate greater structural confidence). While a trade-off in scRMSD was observed compared to RFdiffusion alone, a substantial proportion of designs still achieved scRMSDs below 3 Å (Fig. 2b).

However, our primary objective was to assess backbone novelty. To this end, we selected backbones with high designability (scRMSD <3 Å and pLDDT >90) and performed structural similarity searches against the CATH database^22,23^ with FoldSeek^24^, reporting the highest template modeling score (Max PDB TM-score^25,26^). By utilizing low-frequency secondary structure descriptor guidance, our approach produced significantly more structurally novel and highly designable backbones compared to baseline RFdiffusion (Fig. 2c). This suggests that DiffTopo-RFdiffusion can generate both native-like folds and previously unobserved (dark) folds, demonstrating generalization beyond the structural space of the Protein Data Bank (PDB).

As an example, for the secondary structure descriptor seed “EEEHHEHE”, we generated 500 designable backbones which resulted in 109 structurally distinct clusters (Fig. 2d), using a TM-score threshold of 0.5^27^.These findings highlight the ability of our framework to efficiently generate a diverse range of *de novo* protein structures with low structural redundancy.

In summary, the DiffTopo-RFdiffusion framework enhances de novo protein design by generating designable and structurally novel backbones. Compared to standalone RFdiffusion, it significantly improves backbone novelty by leveraging sampling in low-frequency secondary structure order regions, enabling the discovery of both native-like and entirely dark folds beyond the known protein space.

### Exploring de novo “dark” folds with DiffTopo

A primary goal of our work is to expand the accessible protein fold space by generating structures that are not only new to nature but also extend beyond the range of folds observed in natural proteins. To systematically assess the model’s capacity for fold exploration, we sampled 779 different dark folds with distinct secondary structure compositions (all-α, all-β, and α/β), and quantified their statistics on the designability, novelty, and diversity (Supplementary Table 1). At the sequence level none of the designed sequences displayed a hit below the significant E-value threshold of 0.001 (Supplementary Table 2) relative to the non-redundant database (NR)^28^, indicating that the designed sequences are novel and distinct from naturally occurring sequences. For additional validation, we applied the HHpred^29,30^ to search for potential distant homologs. Intriguingly, all sequences showed E-values above 0.5, further supporting their novelty compared to known natural sequences (Supplementary Table 3).

To further evaluate the structural quality and novelty of the generated designs, we analyzed sequence-structure consistency, architectural diversity, and similarity to known folds (Fig. 2e–h). As shown in Fig. 2e, DiffTopo produced backbones with high designability across a range of secondary structure compositions, indicating strong sequence-structure compatibility. To illustrate this, we selected representative examples of all-α, all-β, and α/β folds and showed their corresponding structures in the figure; corresponding secondary structure descriptors are provided in Supplementary Table 4. The resulting protein sketches exhibited substantial architectural diversity (Fig. 2f), showing that the model explores a wide topological space. Structural alignment with AF2 predictions confirmed the internal consistency of these designs (Fig. 2g), supporting their plausibility. The designs were compared to naturally occurring structural databases using Foldseek, in which the closest hits had very low structural similarity, underscoring the potential of the approach for discovering novel, previously uncharacterized folds (Fig. 2h).

To evaluate the novelty of the generated folds, we compared all dark folds to CATH domain representatives using TM-align and projected the TM-score matrix into 2D using classical multidimensional scaling (MDS) (Supplementary Fig. 2). The resulting distribution showed that dark folds form distinct structural clusters, separate from natural domain families. Even for well-populated classes like 7-helix bundles, with 394 CATH entries, DiffTopo still generated 16 unique topologies (Max TMscore <0.5) not accounted for in the database (Supplementary Fig. 3, Supplementary Table 5). To further explore their global distribution in fold space, we applied the latest structural analysis method SHAPES^31^ (structural and hierarchical assessment of proteins with embedding similarity), projecting structures into two orthogonal spaces, ProtDomainSegmentor embedding^32^ and ProteinMPNN layer 1 embedding (see Methods: Mapping Protein structures to CATH database). As shown in Supplementary Fig. 4, each embedding space was rasterized into a 16×16 grid, and representative structures were selected per cell to visualize sampling coverage. While CATH domains densely populated specific regions, the dark folds broadly covered previously unoccupied areas, supporting that our method systematically samples novel and underexplored fold topologies.

In summary, our pipeline demonstrates its ability to uncover novel, designable and complex folds that do not exist in nature, encompassing various secondary structure combinations, including all *α*, all *β*, and mixed *α*/*β* folds.

### DiffTopo dark folds designs exhibit high structural precision

We computationally designed 778 dark folds that passed the designability filters. For experimental validation, we selected 12 all-α, 7 all-β, and 21 α/β dark fold designs. These proteins were expressed in *Escherichia coli* and purified using a Ni^2^+-NTA affinity column. The purified proteins were characterized by size-exclusion chromatography combined with multi-angle light scattering (SEC–MALS) and circular dichroism (CD) spectroscopy. Of 44 tested dark fold topologies, 22 designed proteins were well expressed and highly soluble. Additionally, all designs exhibited CD spectra typical of folded proteins and consistent with the designed secondary structure content (Supplementary Fig. 6-10). The designs were thermally very stable as is typical in de novo designed proteins, and were confirmed to be monomeric in solution by SEC– MALS (Supplementary Fig. 5 a-w).

From this set, we selected 6 representative dark folds, including all-α (N34), all-β (N40), and α/β (N5, N10, N30, N37), which are highlighted in Fig. 3a. Crystal structures were solved for 3 of these designs: N30, N37, and N5. For N30 and N37, the crystal structures aligned especially well with the computational models, yielding backbone RMSDs of 0.58 Å and 0.70 Å, respectively (Fig. 3b,c). FoldSeek analysis confirmed minimal similarity to known protein structures (Foldseek CATH max TM-score = 0.488 in), validating their structural novelty. For example, while N37 partially overlapped with ribonuclease H-like β-sheets (Foldseek CATH max TM-score = 0.55 for shared region), its overall topology, α-helices wrapping a β-sheet, is absent from the natural repertoire. Notably, N5, an α/β dark fold (max TM-score = 0.43), crystallized as a dimer with a β-strand-swapped interface, however, SEC–MALS confirmed its monomeric state in solution, and alignment of the non-swapped regions (35-161) revealed strong agreement with the design model (full atom-RMSD = 0.66 Å), suggesting that the strand swapping event is likely a crystallization induced artefact.

**Figure 3.**
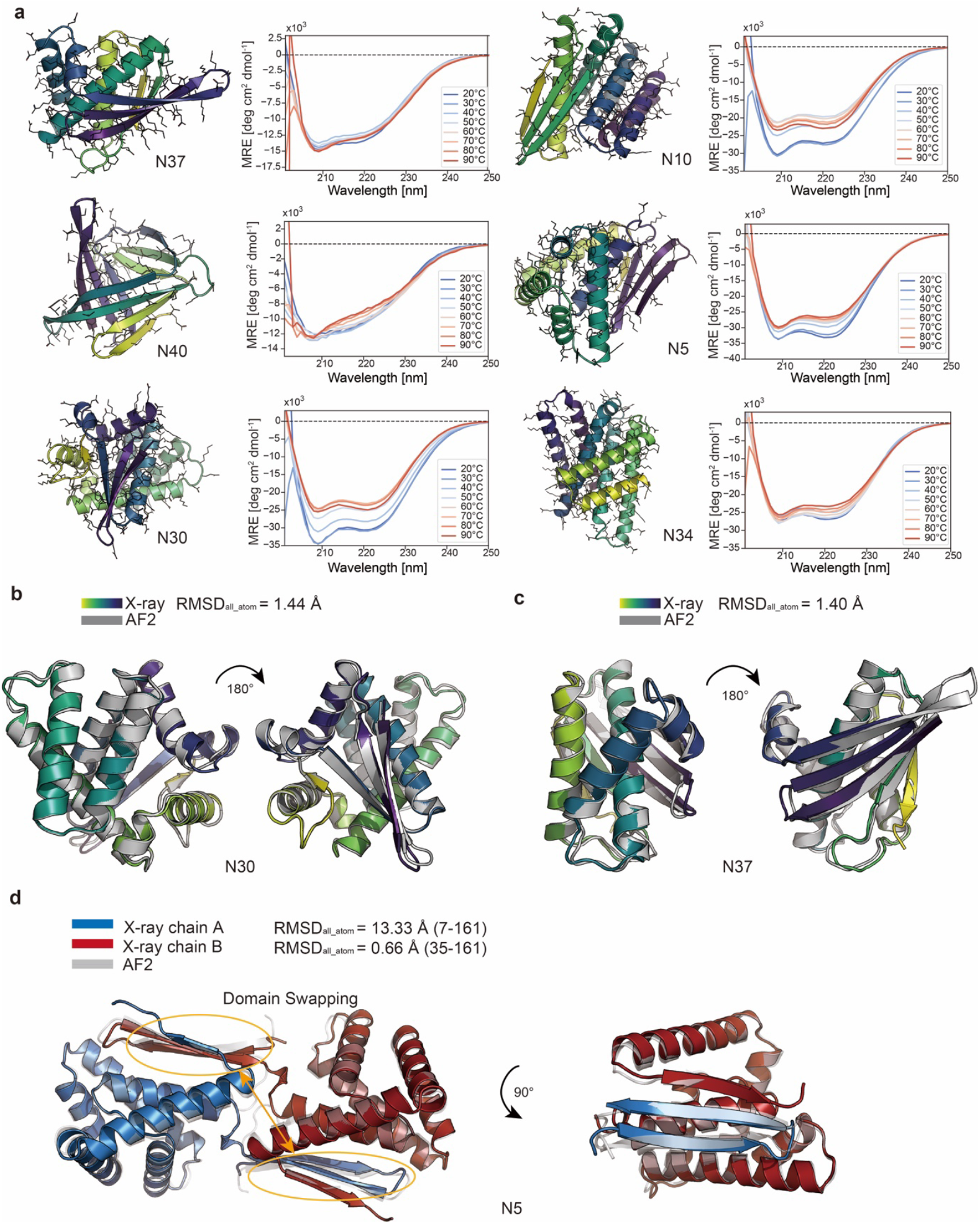
Experimental validation of dark folds. **a**, structural models of representative designs from each fold category and their corresponding circular dichroism (CD) spectra at different temperatures, confirming that the designs adopt folded structures in solution. **b**, X-ray crystal structure of N30 (colored) superimposed on the AF2 predicted model (gray). **c**, X-ray crystal structure of N37 (colored) superimposed on the design model (gray). **d**, X-ray crystal structure of N5 chain A (red) and chain B(blue) overlaid with the AF2 predicted model (gray), shown as a dimer (left) and a monomer (right). Crystal structures are available in PDB: N30(9R2K), N37(9R2L), N5(9R2O).

These results demonstrate that dark fold designs generated by DiffTopo are not only highly designable and diverse, but also structurally accurate, validating the potential of our topology-first framework to generate entirely novel folds absent from nature.

### MirrorTopo generates new to nature protein folds

Although DiffTopo enables the generation of novel folds, the sampled structures still follow natural-like spatial organization. To further expand the design space, we developed MirrorTopo, which generates symmetry-inverted (mirrored) versions of natural folds at the topological level, producing structures that are physically valid but have not been observed in current structural databases.

MirrorTopo operates by mirroring CG topology sketches in 3D coordinate space prior to atomic-level generation using RFdiffusion. Mirror folds exhibit distinct pairwise patterns of secondary structure stacking, specifically in the dihedral angles and distances between helix–helix, helix–strand, and strand–strand pairs compared to natural proteins (Supplementary Fig. 1d,). These patterns fall outside the typical distribution learned by DiffTopo, suggesting that such topologies are unlikely to be sampled. Unlike previous research^33,34^ that mirrored atomic-level coordinates using D-amino acids and right-handed helices, MirrorTopo maintains L-amino acids and right-handed helices, but systematically inverts how secondary structure elements are arranged in space (Fig. 4a). This approach enables the sampling of mirrored topologies that DiffTopo cannot access allowing us to explore fold topologies that are physically valid yet absent from the known natural repertoire.

**Figure 4.**
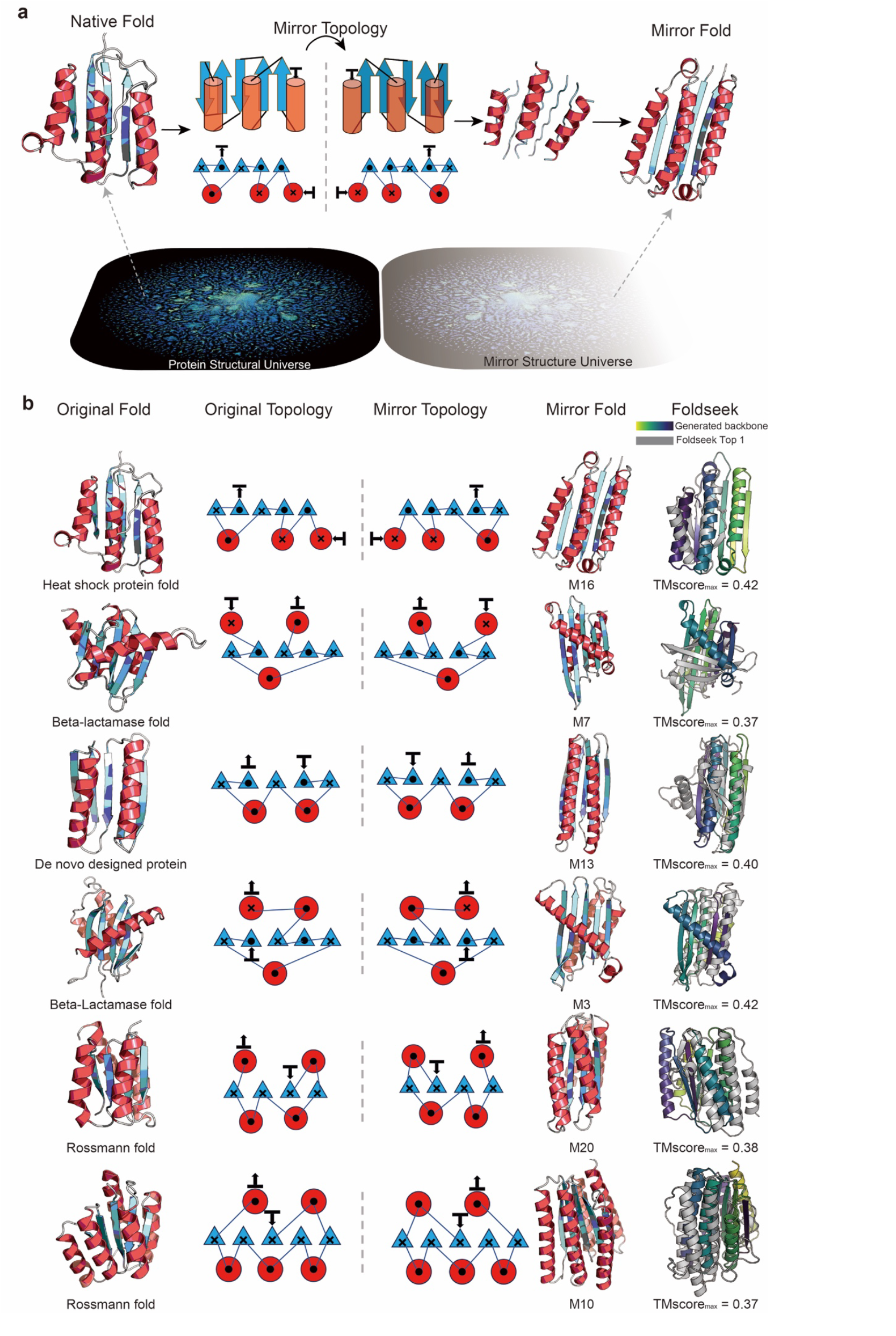
Structural exploration in the Mirror fold space. **a**, Schematic overview of the MirrorTopo pipeline. Native CG topologies are mirrored in the topological coordinate space to generate symmetry-inverted representations. These are then processed through the same downstream design pipeline, including RFdiffusion for backbone generation. The bottom row depicts the natural protein universe derived and mirror fold universe. **b**, From left to right: native protein structure from the CATH database; 2D secondary-structure connectivity of the native fold; mirrored topology reflecting a symmetry-inverted design; backbone model generated from the mirrored topology using RFdiffusion; and the top FoldSeek match from the PDB database, indicating similarity or novelty relative to known folds.

First, we selected 8 native fold types from the CATH database and generated their mirrored topologies using our MirrorTopo pipeline. For each fold, multiple designs passed our designability filters (self-consistency RMSD < 3 Å and AF2 pLDDT > 90). We then selected 21 of these designs for experimental testing. Among them, 13 designs were successfully expressed in *E*.*-coli* and showed expected physicochemical properties, indicating folded proteins. These 13 designs cover 6 different mirror folds from native proteins, spanning a variety of α/β architectures, representing diverse structural classes. All detailed description and structural annotations and CATH classifications are summarized in Supplementary Table 6. The first column of Fig. 4b shows native protein structures from the CATH database, followed by their 2D secondary-structure schematics in the second column. Their mirrored counterparts, obtained by flipping the arrangement of secondary-structure elements, are presented in the third column, with the corresponding backbone models generated by MirrorTopo shown in the fourth column. The rightmost column displays FoldSeek search results, confirming that the mirror folds are structurally novel with no close matches in existing structural databases.

### Mirrored topologies show high structural precision

For the 6 different mirror folds in Fig. 4b, we selected one representative design from each, all of which showed CD spectra consistent with the expected secondary structure content (Fig. 5a) and SEC–MALS profiles indicative of monomeric states in solution (Supplementary Fig. 11 a-l). CD spectra and melting temperature profiles for the remaining successful designs are provided in Supplementary Figs. 12-14. We successfully solved the crystal structures of two mirror fold designs, M16 and M7. M16, the mirror version of the histidine kinase-like ATPase fold, exhibited remarkable agreement with its AF2 prediction (Cα RMSD = 0.42 Å; all-atom RMSD = 1.50 Å), confirming the high structural accuracy of the design (Fig. 5b). Similarly, M7, a mirror of the TRAPP domain (β-lactamase-like topology), also closely matched its computational model (Cα RMSD = 0.67 Å; all-atom RMSD = 1.50 Å) (Fig. 5c). These results validate that mirror topologies, that although absent from nature, can be realized with high fidelity using computational methods and experimentally validated.

**Figure 5.**
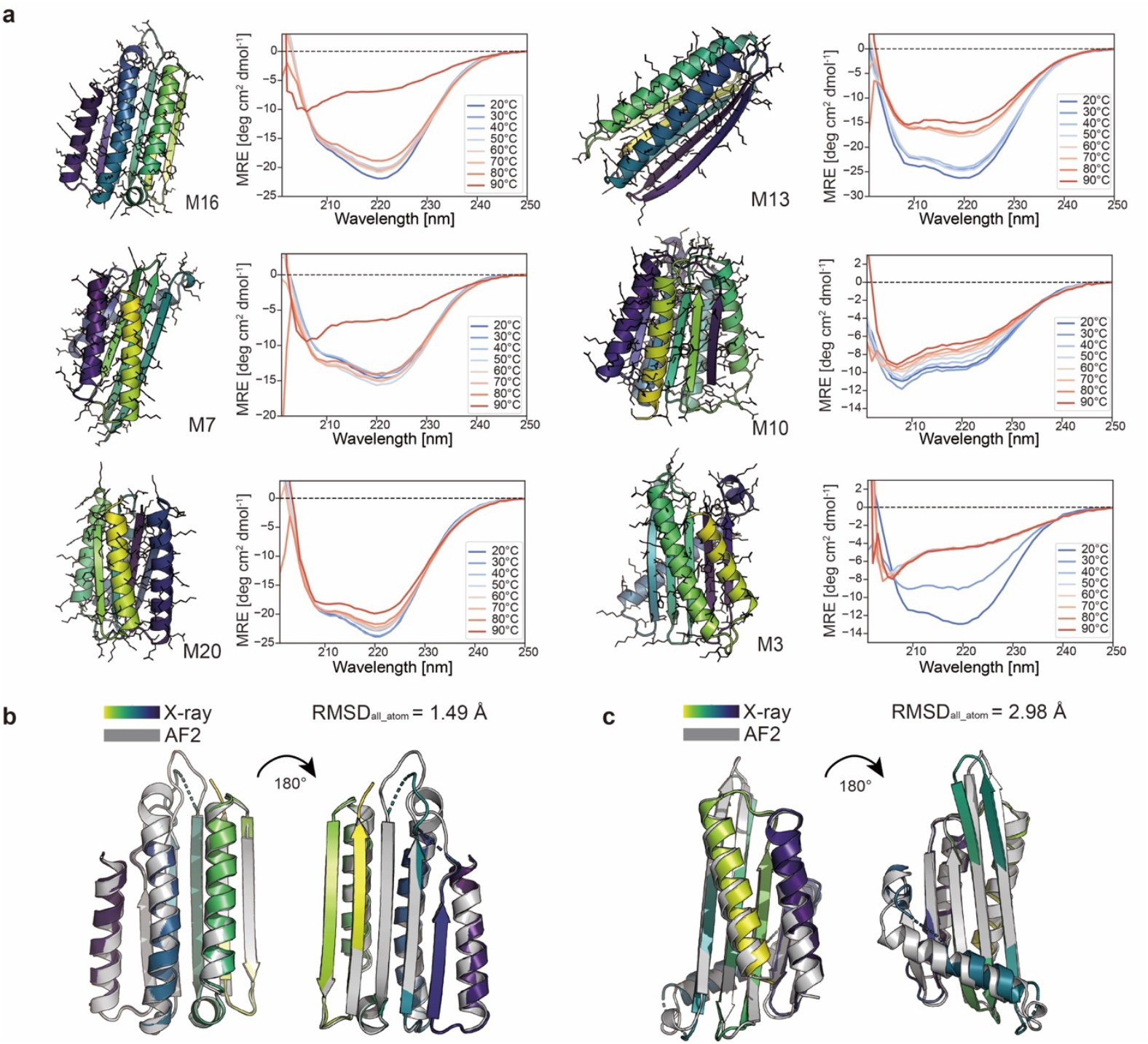
Experimental validation of the computational designed Mirror folds. **a**, Structural models of representative designs from each fold category and their corresponding circular dichroism (CD) spectra at different temperatures, confirming that the designs adopt folded structures in solution. **b**, X-ray crystal structure of M16 (colored) superimposed on the design model (gray). **c**, X-ray crystal structure of M7 (colored) superimposed on the design model (gray). Crystal structures are available in PDB: M16(9R2V), M7(9R2R).

To evaluate whether the tertiary structure motifs in our mirror folds are also found in natural proteins, we used TERMANAL^35,36^. According to previous work by Mackenzie et al., about 600 tertiary structural motifs (TERMs) are sufficient to describe 50% of structures in the PDB. TERMANAL analysis returns two main results: Abundance and Structure Score. Abundance reflects how frequently a particular TERM (tertiary structural module) appears across the protein structure database, with higher values indicating that the TERM is common and likely favorable in natural proteins, whereas lower values indicate rarity and potential structural novelty. Structure Score provides a global assessment of a protein structure, integrating the abundance of all its constituent TERMs to measure overall structural reasonableness. Here, we performed TERM analysis on 50 random CATH folds, all experimentally generated dark folds, and all mirror folds. As shown in Fig. 6a, CATH folds exhibit high motif abundance, with an average of ~0.8 per TERM. Dark folds have moderately lower abundance (~0.68), while mirror folds display the lowest values (~0.44). A similar trend was observed in Structure Scores. To illustrate these results at the residue level, we selected two representative dark folds (N30 and N37) and two mirror folds (M16 and M7), as shown in Fig. 6b,c.The results showed that mirror folds (M16 and M7) have low motif abundance and low structural similarity scores across all regions, suggesting that their local structures are distinct from those commonly found in nature. In contrast, the dark folds N30 and N37 had high motif abundance and structural scores, especially N37, which was similar to native-like topologies. N30 also showed strong matches, except for its β-sheet region, which had fewer matching motifs. These results support our previous observation that DiffTopo samples structures that remain within a “natural-like” distribution, while mirror folds generated by MirrorTopo lie in a different and less explored region of the structural space. In other words, mirror folds access areas of the fold space that are not well covered by natural evolution.

**Figure 6.**
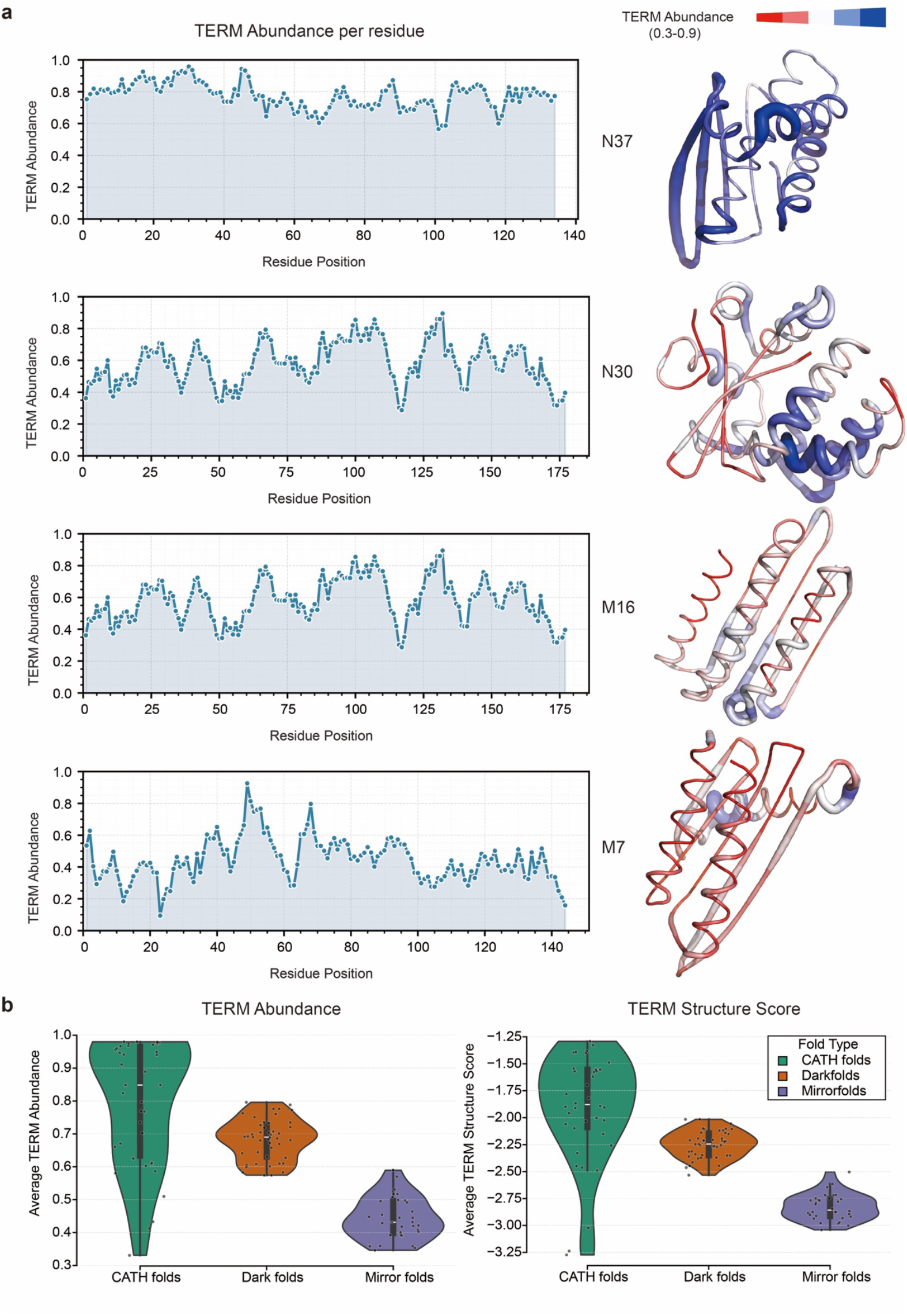
TERM analysis of dark folds and mirror folds. **a**, TERM abundance per residue(left) and representative PDB structures colored by TERM abundance, red-thin regions indicating low abundance; blue-thick regions reflecting higher abundance (right). **b**, Average abundance distribution (left) and structural scores(right) for CATH folds, dark folds, and mirror folds.

Both computational results and experimental validation demonstrate that the MirrorTopo pipeline is viable for generating structurally precise, stable, and novel proteins. The success of mirror-topology designs achieved using right-handed helices with canonical L-amino acids challenges conventional assumptions about foldability and suggests the existence of a vast, underexplored mirror-fold space. These findings highlight the potential of mirror topologies as a complementary approach to expand the universe of designable protein structures.

## Discussion

Exploring and expanding the protein fold space remains an important challenge in protein design, particularly given that new structures could, in principle, enable novel functional roles for proteins. Existing generative models tend to stay close to natural-like topologies due to fine-grained representations and biases in training data. In contrast, our method uses a compact coarse-grained topology representation, that captures only the relative positions and connections of secondary structure elements. With this abstraction, DiffTopo efficiently learns spatial patterns and samples diverse and novel fold topologies beyond those seen in nature. Embedding analysis (Supplementary Fig. 4) shows that many designs occupy underexplored regions of fold space. Despite their novelty, these structures still retain high designability, allowing for sequences that are in good agreement with the structural templates resulting in structural predictions with high AF2 confidence. These results suggest a large, untapped fold space that can now be explored systematically.

While DiffTopo expands the diversity of folds, it still follows the spatial patterns found in natural proteins. To move further beyond these constraints, we introduced MirrorTopo, which generates mirrored topologies that are structurally valid but evolutionarily absent. These mirror folds use standard L-amino acids and form right-handed helices, yet their global secondary structure arrangement is reversed. They do not resemble natural folds, as confirmed by FoldSeek analysis, and occupy regions rarely reached by generative models. Embedding analysis shows these mirror designs fall far outside the distribution of natural structures (Supplementary Fig. 1d), suggesting that mirror fold space represents a distinct, orthogonal region of protein architecture.

The absence of mirror folds in nature could reflect evolutionary selection rather than physical constraints. Many natural folds have emerged through duplication, fusion, and recombination events that favor certain packing orientations. Mirrored architectures, while physically allowed, may be disfavored due to folding kinetics, sequence-to-structure mapping biases, or evolutionary path dependence. By systematically constructing these mirror topologies, MirrorTopo reveals a vast, underexplored region of the fold space that has remained invisible to both evolution and previous generative methods. This space is not only accessible, but it is designable, stable, and experimentally verifiable.

To test whether these topologies are physically viable, we experimentally solved five crystal structures across both DiffTopo and MirrorTopo designs. These include dark folds (N30, N37, N5) and mirror folds (M16, M7). The results confirmed that these unconventional topologies can fold precisely into their intended atomic structures, supporting the robustness of our design approach.

Among all dark folds, N30 displays a particularly entangled topology, while most other designs rely on linear stacking of secondary structures. In its crystal structure, one helix (residues 120–143) threads through a closed cavity formed by two others (61– 76 and 81–95), creating a topological constraint unlikely to arise via simple stacking. Sequential AF2 prediction analysis suggests a nonlinear folding pathway: the first two helices fold and pack tightly, and only later open to accommodate the third helix (Supplementary Figure 15 and Supplementary information: Sequential AF2 predictions analysis for N30). This behavior expands the known repertoire of foldable structures and suggests that topological complexity—such as threading or knot-like arrangements—can be encoded and accessed through designed sequences. N30 highlights the potential of our framework to generate not only novel folds, but also nontrivial folding mechanisms.

While this work primarily focuses on structural diversity, our method also supports functional design through motif scaffolding (see Methods: Motif-Scaffolding via Conditional Inpainting with Diffusion Models). By conditioning on existing secondary structure motifs, DiffTopo can also generate diverse scaffolds that preserve motif geometry), highlighting the potential of our framework to support functional scaffold generation, expanding the possibilities for designing proteins with tailored interactions. Overall, our study shows the potential of utilizing different representations to enhance the exploration of ML-based methods for generative protein design showing that under sampled regions of this landscape can be accessed through this approach. We anticipate that these new methods will in future enable the design of proteins with novel biological functions.

## Methods

### Coarse-grained topology representation

Inspired by the FORM representation^12^ and protein sketches^13,14^, here protein structures are built by the assembly of their secondary structures, we adopt a simplified representation referred to as a “coarse-grained topology (CG topology)”. The topology reduces the intricate protein structure into a stacked arrangement of Secondary Structure Elements (SSEs), achieved by representing secondary structures with three carbon alpha centroids, as illustrated in Fig. 1a. All the secondary structures in each residue number are assigned by STRIDE^37^ program. CG topology involves capturing the geometric position of helices and strands through centroids. For helices, centroids include the first and last four C*α* atoms, as well as the total C*α* atom centroid. Strands are represented by centroids of the first and last two C*α* atoms, along with the centroid C *α* atom within the strand. CG topology can also be easily represented as a protein sketch, which is a rough 3D approximation of a native protein structure with standard SSEs, lacking loops, and AA side chains. Compared with forms and protein sketches, this representation method of CG topology removes the concept of layers, offering a higher degree of freedom in structural representation to represent structures such as beta barrels, while remaining simple with few degrees of freedom.

### Dataset

Using the CG topology approach we can easily convert a protein structure database, such as PDB, from standard protein models to CG topology structures. In this paper, all PDB structural data are sourced from CATH^22^, a database organized as a classification of protein structures. To mitigate the influence of structural redundancy on the data distribution, we employ the CATH-dataset-nonredundant S40 for training data, yielding 31,886 non-redundant protein structures. 90% of structures are used for training and 10% are used for independent validation.

### DiffTopo/MirrorTopo + RFdiffusion framework

First, we introduce DiffTopo, a Equivariant diffusion model^38,39^ for generating CG topology conditioned on SSEs strings. Based on previous works on denoising diffusion, given a data point sampled from a real data distribution *z*_0_ ~ *q*(*z*), the forward diffusion process is defined by adding Gaussian noise gradually to the data point *z*_0_ from corrupted samples *z*_*t*_. For one CG topology data *z*_0_, there are three points *x* and corresponding secondary structure type *u*. At time step *t* = 0, …, *T*, the conditional distribution of the intermediate data state *z*_*t*_ given the previous state is defined by the multivariate normal distribution,

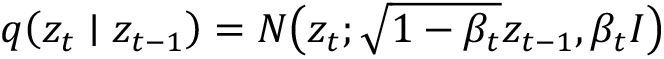

The step sizes are controlled by a variance schedule 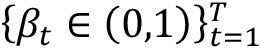. The process is constructed to be Markovian and if we let *α*_*t*_ = 1 − *β*_*t*_ and 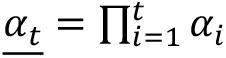, we can obtain the distribution of *z*_*t*_ given *z*

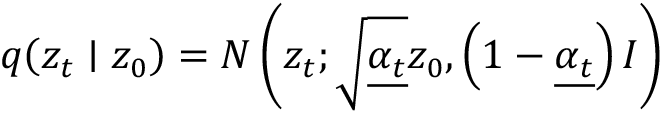

For the backward process, we need to learn a model *p* to approximate these conditional probabilities.

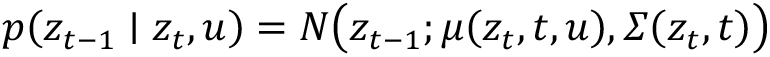

*μ*_*θ*_(*z*_*t*_, *t, u*) is predicted by the neural network. In this paper function *μ* that predicts noise *ϵ*_*θ*_ is implemented as geometric vector perceptron (GVP)^40^. The input to GVP is the noised version of the point coordinates *z*_*t*_ and point feature *h*_*t*_ at time *t* and context *u*. Note that the predicted noise *ϵ* includes coordinate and feature components, *ϵ* = [*ϵ*_*z*_, *ϵ*_*h*_]. The predicted noise can be calculated by

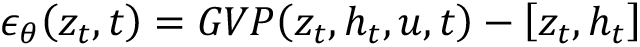

In addition to DiffTopo, we introduce MirrorTopo, a method for generating mirror-image topologies directly from native protein folds. Given a CG representation of the native fold, we obtain the mirror topology by inverting the z-coordinates while preserving the x- and y-coordinates. This transformation effectively reflects the structure across the xy-plane, yielding a topologically mirrored conformation without altering sequence or connectivity.

Following the generation of the coarse-grained (CG) topology via the diffusion process or mirroring of the native CG topology, the resulting topology enables the direct extraction of the positions and lengths of standard SSEs. Since CG topology retains a certain level of fuzziness, we need a backbone level model to systematically search for the designable backbone corresponding to CG topology. Here, we leverage RFdiffusion, a highly effective backbone generation model to generate designable backbones from CG topology. Initially, we construct a protein sketch from standard SSEs based on CG topology and utilize the motif modeling function inherent in RFdiffusion to directly connect the SSEs. Subsequently, we employ the partial diffusion approach to identify a reasonable backbone structure within this interconnected structure. Then we employ the ProteinMPNN^18^ algorithm for fixed backbone sequence design to obtain amino acid sequences. These sequences are then input for the structure prediction algorithm AF2^19^ to predict the structure, which is use to validate the designability of generated backbones.

### Secondary structure descriptors

#### Secondary structure descriptor construction

To define secondary structure conditions within a specific topological class, we constructed strings representing the desired sequence of secondary structure elements—for example, “EEEHHEHE”, where each character denotes a secondary structure type (E: β-strand, H: α-helix). These SSE strings were generated either manually or through algorithmic enumeration by specifying the number, type, and order of elements. The example “EEEHHEHE” encodes a topology with eight secondary structure elements—five β-strands and three α-helices arranged in a particular sequence. These strings were used as input constraints to the generative model, directing the synthesis of backbones consistent with the specified topology.

#### Random Secondary Structure Descriptor Sampling

We randomly selected 512 unique SSE order strings, ranging in length from 3 to 12 elements, from the CATH database. These sequences served as topological constraints to guide the generation of varied backbone conformations.

#### Low-Frequency Secondary Structure Descriptor Sampling

To specifically target rare topologies, we compiled all possible SSE order strings of length 8 and computed their frequency across the entire CATH database. Strings with fewer than two occurrences were classified as low-frequency compositions. This procedure yielded a total of 2,471 underrepresented SSE patterns, which were used to probe the model’s ability to generalize beyond commonly observed folds.

### Mapping Protein structures to CATH database

We computed ProteinMPNN encoder and ProtDomainSegmentor^22^ embeddings for all CATH structures from Ingraham et al.^29^, selecting only those without chain breaks, with resolution better than 3.0 Å, and R_free_ below 0.25. Both embedding types are computed at the per-residue level and averaged along the sequence dimension to obtain 128-dimensional embeddings for ProteinMPNN and 4096-dimensional embeddings for ProtDomainSegmentor. To capture structural features at different hierarchical scales, we extract local fold features using the first encoder layer of ProteinMPNN, which encodes featured geometries of nearest neighbors, while global fold features are captured using the pre-final layer of ProtDomainSegmentor, which is predictive of CATH Architecture classification. We project the CATH embeddings onto their first two principal components (PCs) and apply the same transformation to embeddings of dark fold and mirror fold structures for direct comparison.

### TERM analysis for protein structures

For TERM analysis, we randomly selected 50 representative protein structures from the CATH database, including both dark folds and mirror folds that have been experimentally expressed. All structures were queried using the TERMANAL software package against the default fragment database “bc-30-sc-20141022”, a non-redundant subset of protein chains in the Protein Data Bank (PDB) with a 30% sequence identity cutoff and determined by X-ray crystallography at high resolution. The resulting abundance and structural score files were used for subsequent analyses.

### Protein expression and purification

DNA sequences corresponding to the amino acid sequence of the designed proteins were optimized for E. Coli codon usage and purchased from Twist Bioscience as fragments, including a solvent exposed C-terminal poly histidine tag. The DNA fragments were cloned via Gibson cloning into a pET11b backbone, and plasmids were validated by sequencing. Expression was conducted in BL21(DE3) strain: 0.5 L cultures were inoculated from an overnight culture in Luria-Bertani medium supplemented with ampicillin (100 *μg*/*mL*) and incubated at 37°C and 220 rpm shaking until reaching an OD600 of 0.6-0.8; the cultures were then further incubated overnight at 18°C and inoculated with 0.5 mM IPTG to induce protein expression. Bacterial cells were harvested by centrifugation and pellets were resuspended in 30 mL lysis buffer (100 mM TRIS, pH 7.5, 500 mM NaCl, 5% glycerol, 1 mg/mL lysozyme, 1 mM PMSF, 4 *μg*/*ml* DNase, 0.5X Cell Lytic lysis reagent), incubated at 4°C on a rotating wheel for 1h, before the lysate was clarified by high-speed centrifugation followed by sterile filtration (0.22 *μm*). The lysate was then applied on a 5 ml His-Trap FF column on an ÄKTA pure system (GE Healthcare), the bound proteins were first washed in a Tris based buffer (50 mM Tris, pH 7.5, 500 mM NaCl, 10 mM imidazole) and then eluted with 500 mM imidazole. Polishing purification step was performed by size exclusion chromatography on a Hiload 16/600 Superdex 75 pg column (GE Healthcare) in PBS buffer (pH 7.4). The peak corresponding to the expected molecular weight was collected and concentrated for analysis.

### Circular dichroism spectroscopy

Far-UV circular dichroism spectra were collected between wavelengths of 200 and 250 nm on a Chirascan™ spectrometer (AppliedPhotophysics) spectrometer in a 1 mm path-length quartz cuvette. Proteins were prepared in PBS at a concentration of 0.3 mg/mL by default. Wavelength spectra were measured with a scanning speed of 20 nm/min and a response time of 0.125 s and each measurement was reference subtracted using a cuvette with PBS. The thermal denaturation curves were collected by measuring the change in ellipticity at 220 nm from 20 to 90°C with 2°C increments. Size-exclusion chromatography combined with multi-angle light scattering monodispersity and oligomeric state of the purified proteins were assessed by multi-angle light scattering, thanks to molecular weight determination in solution. 100 *μg* of protein were injected into a Superdex 75 10/300 GL column (GE Healthcare) in PBS buffer (pH 7.4) at a flow rate of 0.5 ml/min and detected on in-line multi-angle light-scattering detector (DAWN TREOS, Wyatt). Static light-scattering signals were recorded, and the scatter data were analyzed by ASTRA software (version 8.0.2.5 64-bit, Wyatt).

### Crystallization and X-ray structure determination

The N5 design was crystallized at a concentration of 19 mg/ml using sitting drop vapor diffusion at 18°C in 0.1 M MES pH 6.5, 0.2 M KSCN, 25% w/v PEG 2000 MME buffer (Clear Strategy Screen I, Molecular Dimensions). The N30 design was crystallized at a concentration of 24.3 mg/ml using sitting drop vapor diffusion at 18°C in 0.1 M MES pH6.5; 30% v/v PEG Smear Low buffer (BCS Screen, Molecular Dimensions). The N37 design was crystallized at a concentration of 28 mg/ml using sitting drop vapor diffusion at 18°C in 0.1 M NaOAc pH5.5, 0.2 M KSCN, 25% w/v PEG 2000 MME buffer (Clear Strategy Screen I, Molecular Dimensions). The M7 design was crystallized at a concentration of 24.7 mg/ml using sitting drop vapor diffusion at 18°C in 0.1 M MES pH6.5, 0.2 M KSCN, 15% w/v PEG 4000 buffer (Clear Strategy Screen I, Molecular Dimensions). The M16 design was crystallized at a concentration of 42 mg/ml using sitting drop vapor diffusion at 18°C in 0.1 M Tris pH8.0, 0.04 M sodium format, 0.04 M CaCl2, 25 % v/v PEG Smear Low buffer (BCS Screen, Molecular Dimensions). Crystals were cryoprotected in 25% glycerol and flash cooled in liquid nitrogen. Diffraction data was collected at the European Synchrotron Radiation Facility MASSIF-1 and MASSIF-3 beamlines, Grenoble, France at a temperature of 100 K. Crystallographic data was processed using the autoPROC package^43^. Phases were obtained by molecular replacement using the full or partial designed model in Phaser^44^. Atomic model rebuilding and refinement was performed using COOT ^45^and Phenix refine^44^. The quality of refined models was assessed using MolProbity^46^.

## Data availability

All data are available in the main text or as Supplementary Information. Atomic coordinates and structure factors of the reported X-ray structures have been deposited in the PDB under accession numbers 9R2K(N30), 9R2O(N5), 9R2L(N37), 9R2R(M7), 9R2V(M16).

## Code availability

DiffTopo code and jupyter notebook is available at GitHub (https://github.com/YangyangMiao/DiffTopo). AF2 model used for \predictions can be downloaded from (https://github.com/sokrypton/ColabFold). ProteinMPNN is available at GitHub (https://github.com/dauparas/ProteinMPNN).

