## Supplementary materials for "Leveraging protein representations to explore uncharted fold spaces with generative models"

### DiffTopo captures Coarse-grained SSE distribution

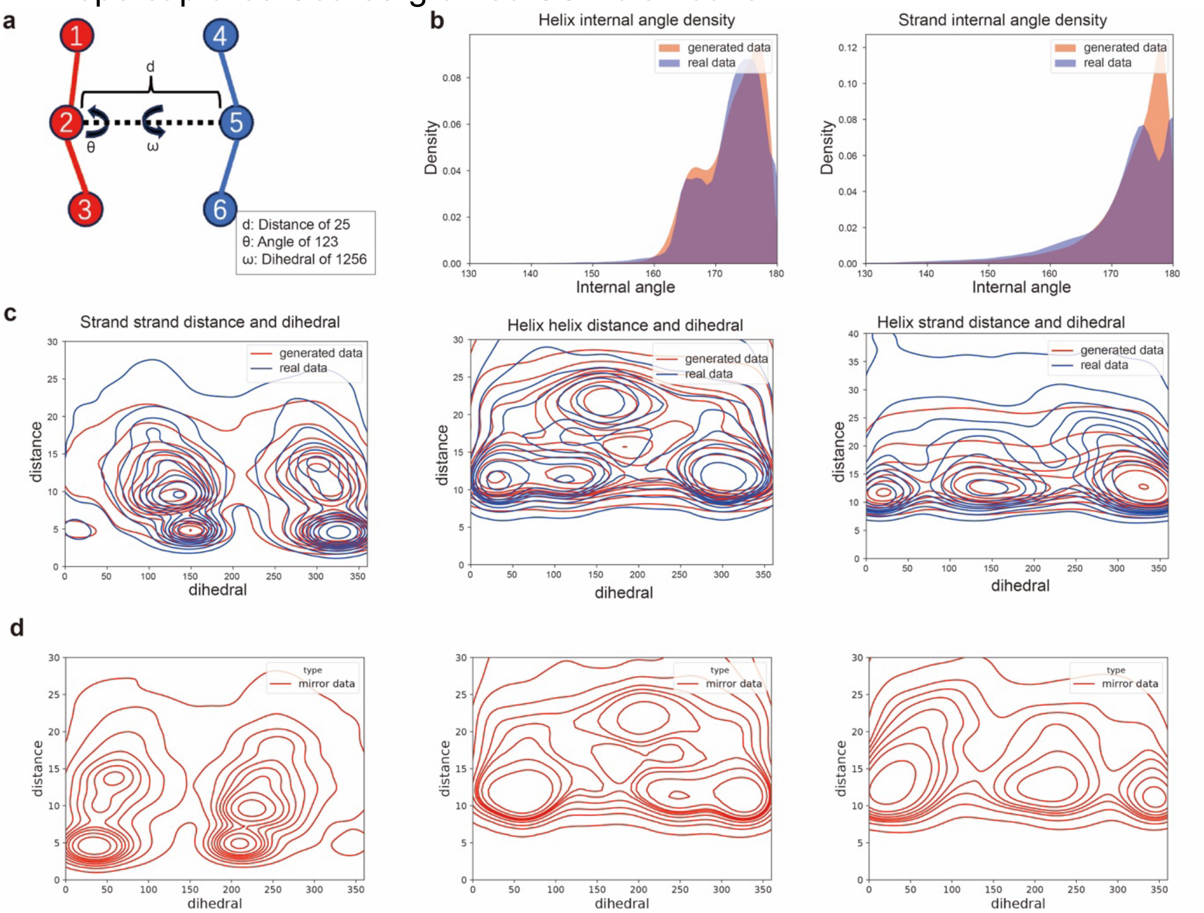


**Supplementary Figure 1. Structural representation and distribution of real and generated data.** **a**, Local and pairwise geometric features to describe spatial relative positions of SSEs, including internal angles within one SSE, distance between centroids of two SSEs and dihedrals between two SSEs. **b**,Local geometric feature distributions of generated (red) and real data(blue). **c**. Pairwise geometric feature distributions of generated (red) and real data (blue). **d**. Pairwise geometric feature distributions of mirrored data (red).

DiffTopo captures the distributions of native protein structures. Standardized metrics are not available to assess the quality of the generated topological sketches. In backbone generation tasks, evaluation often relies on geometric features like Ramachandran distribution and bond length, angle. Based on the sketch level representation, our evaluation focuses on the diffusion model’s ability to estimate data probability density, checking alignment with authentic data distribution. We defined several geometric descriptors for the topological features of (Supplementary Fig1 a). The DiffTopo generated CG topologies yield a similar distribution to the features observed in natural structures (Supplementary Fig1 b,c). The results indicate that the diffusion model adequately approximates the distribution of real data, despite with a tendency to smoothing discrete high-density peaks. While in Supplementary Fig1 d shows the distribution of the mirror CG topologies are in the other side of the space.

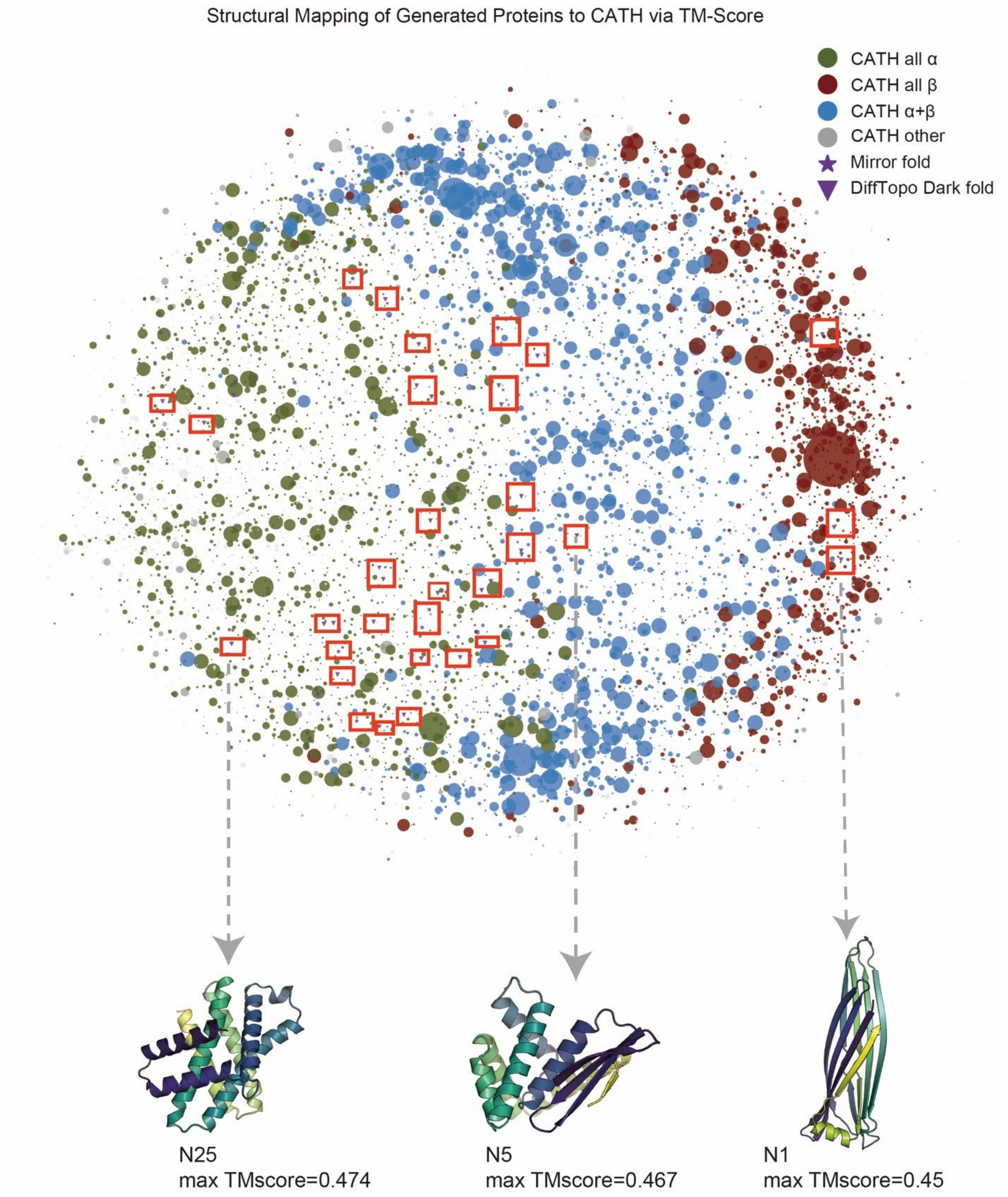


**Supplementary Figure 2. Protein fold space comparison: TM-score-based mapping of designed folds against known structures from CATH, visualized using multidimensional scaling (MDS).** Dark folds are marked with stars, mirror folds with triangles, and native folds are also indicated with circles. Circle size represents the number of members in each fold superfamily. We use red block to mark those split clusters of novel fold and also select 3 represented folds in the bottom .

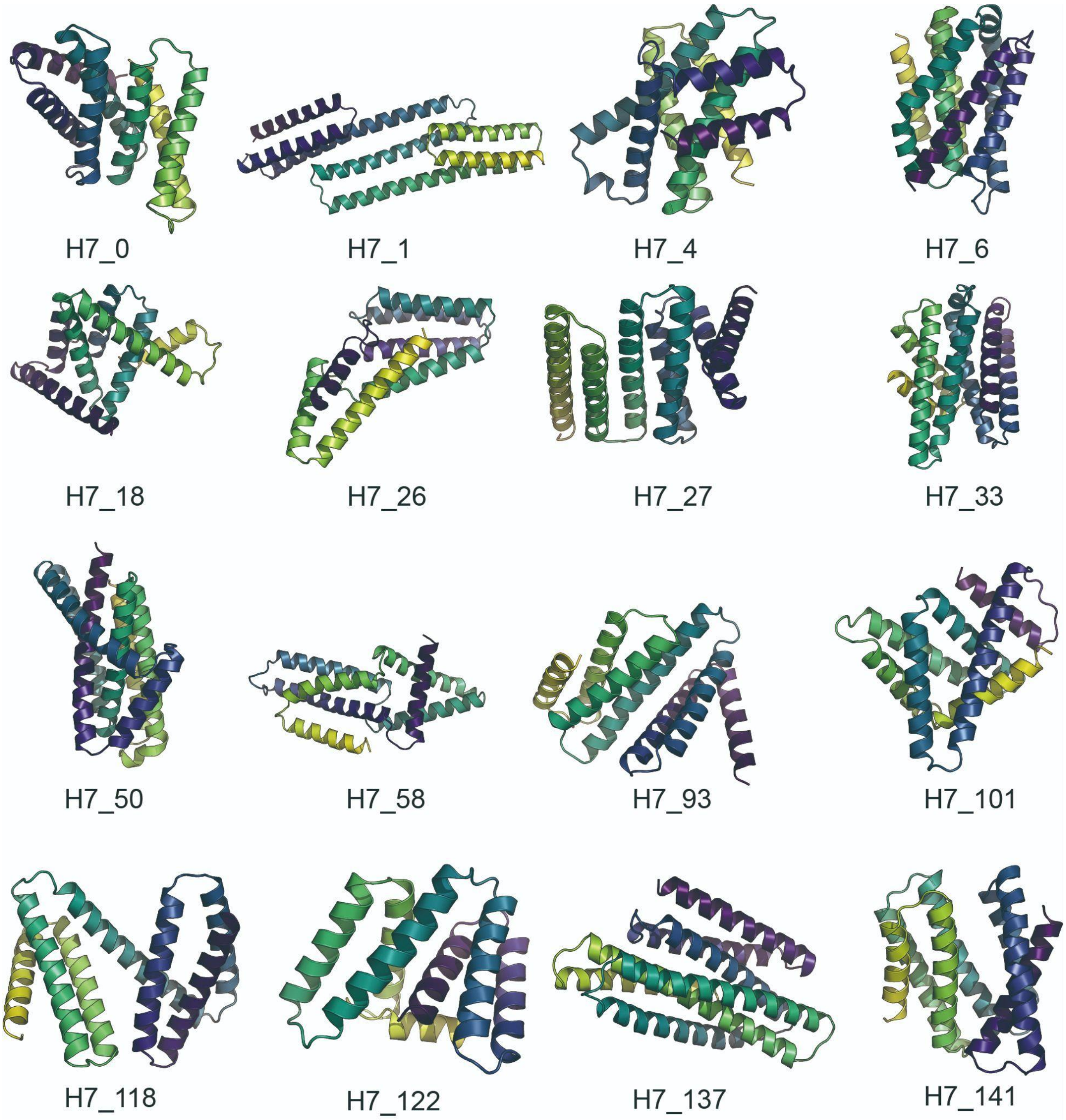


**Supplementary Figure 3. Structural visualization of 16 different novel 7 helix folds**

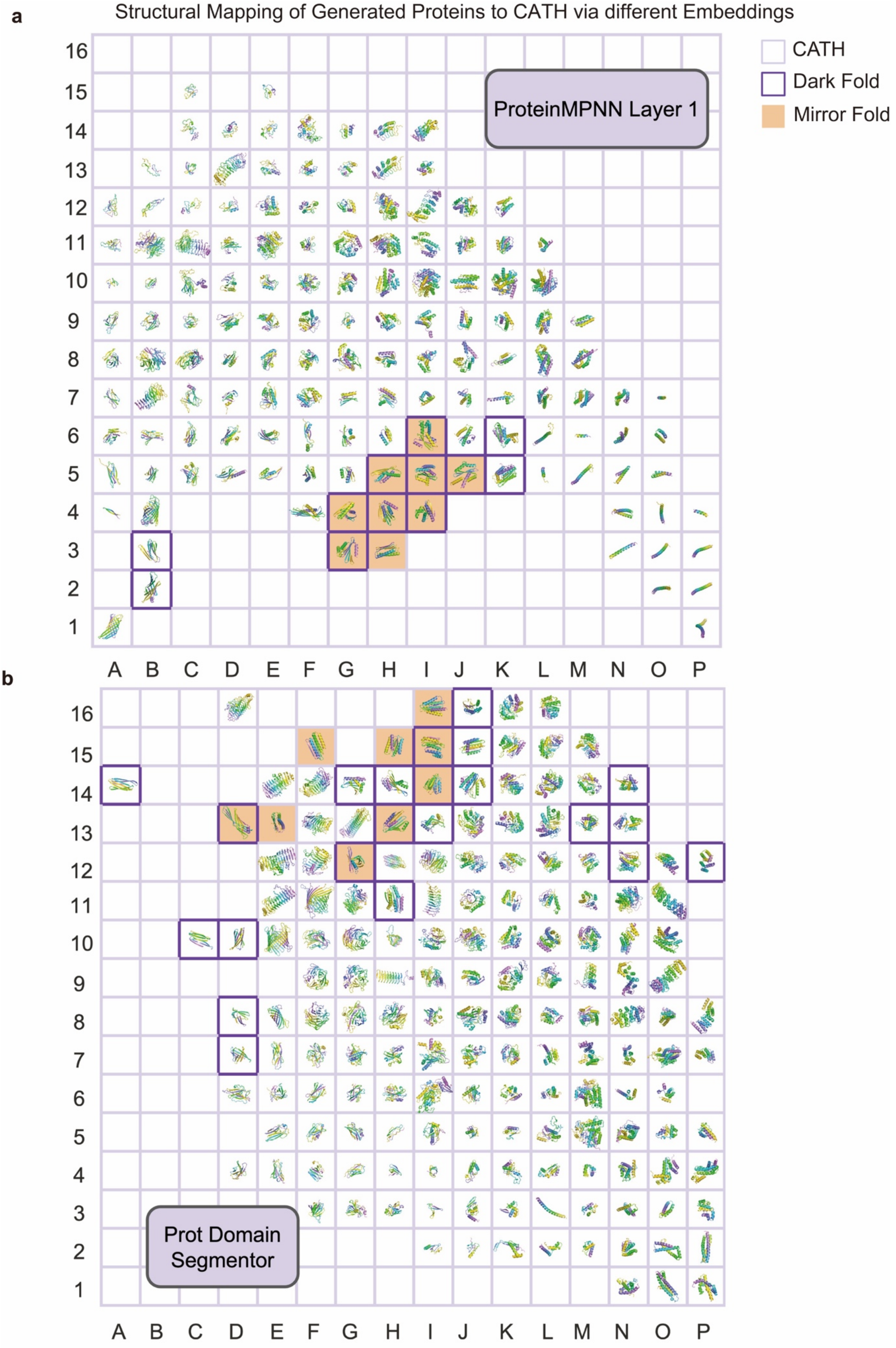
a
Structural Mapping of Generated Proteins to CATH via different Embeddings
CATH
Dark Fold
Mirror Fold
ProteinMPNN Layer 1
16
15
14
13
12
11
10
9
8
7
6
5
4
3
2
1
A B C D E F G H I J K L M N O P
b
Prot Domain Segmentor
16
15
14
13
12
11
10
9
8
7
6
5
4
3
2
1
A B C D E F G H I J K L M N O P

**Supplementary Figure 4. Mapping Generated Structure Embeddings to the CATH Structure Embedding Space.** a, ProteinMPNN Layer 1 embeddings and b, ProtDomain Segmentor embeddings are shown. Each grid corresponds to a representative CATH structure mapped into the 2D grid space. Empty cells indicate regions where no structures are present in the dataset. Notably, purple and orange regions mark embedding areas that lack known native folds in CATH but are populated by our generated designs. These include dark folds(purple) and mirror folds(orange), illustrating that the models explore topological regions beyond the natural protein structure space.

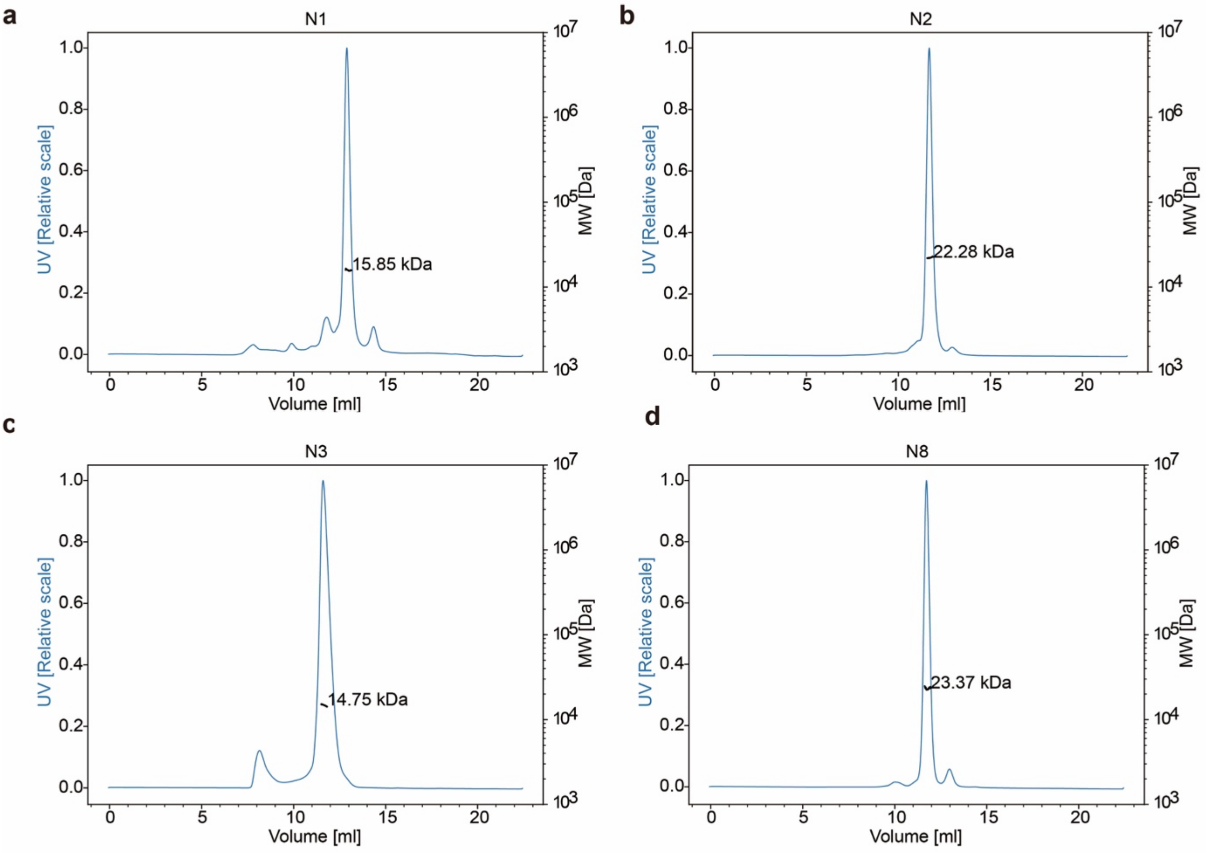

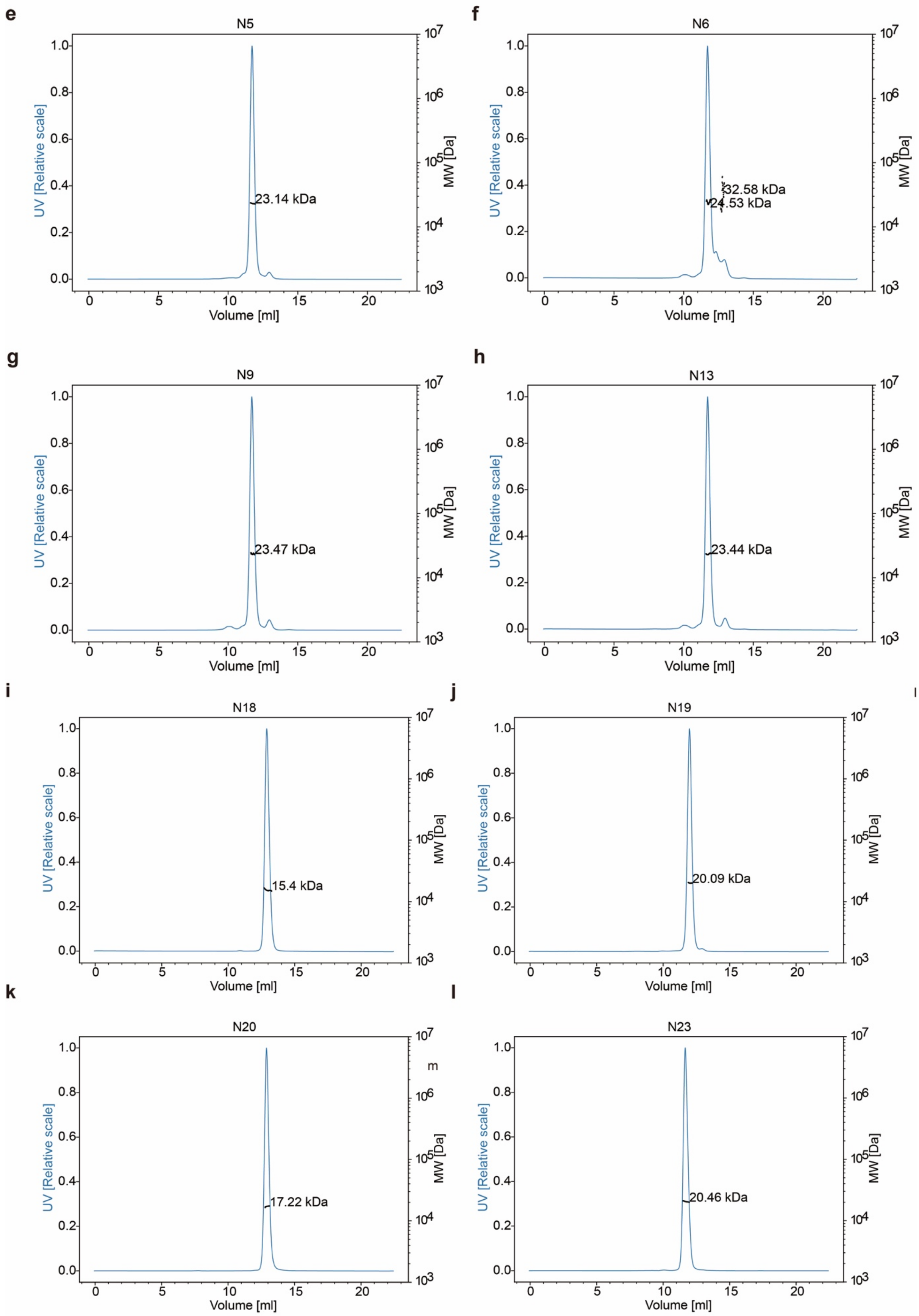
e
N5
23.14 kDa
f
N6
32.58 kDa
24.53 kDa
g
N9
23.47 kDa
h
N13
23.44 kDa
i
N18
15.4 kDa
j
N19
20.09 kDa
k
N20
17.22 kDa
l
N23
20.46 kDa
UV [Relative scale]
MW [Da]
Volume [ml]

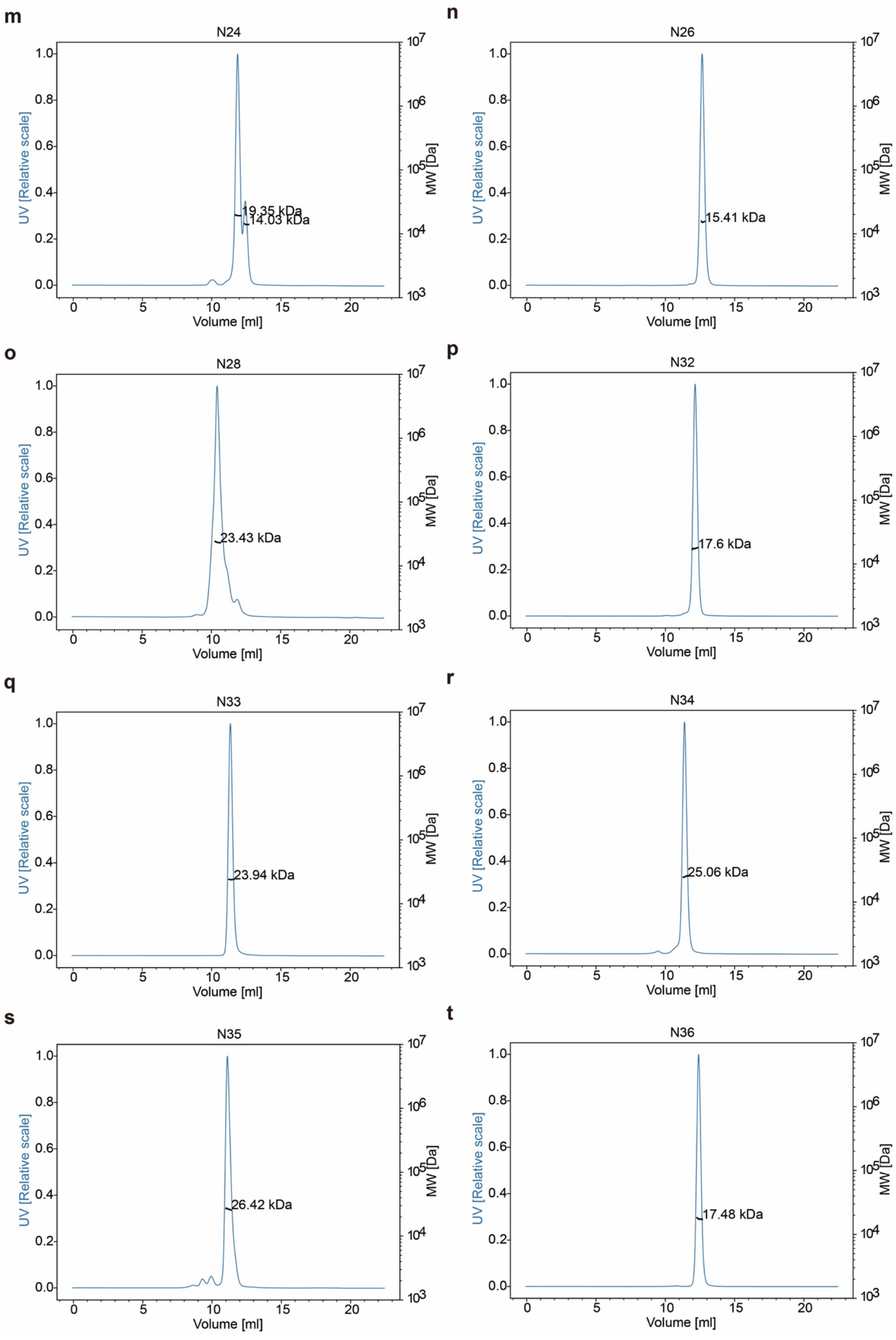

m
N24
19.35 kDa
14.03 kDa
n
N26
15.41 kDa
o
N28
23.43 kDa
p
N32
17.6 kDa
q
N33
23.94 kDa
r
N34
25.06 kDa
s
N35
26.42 kDa
t
N36
17.48 kDa
UV [Relative scale]
MW [Da]
Volume [ml]

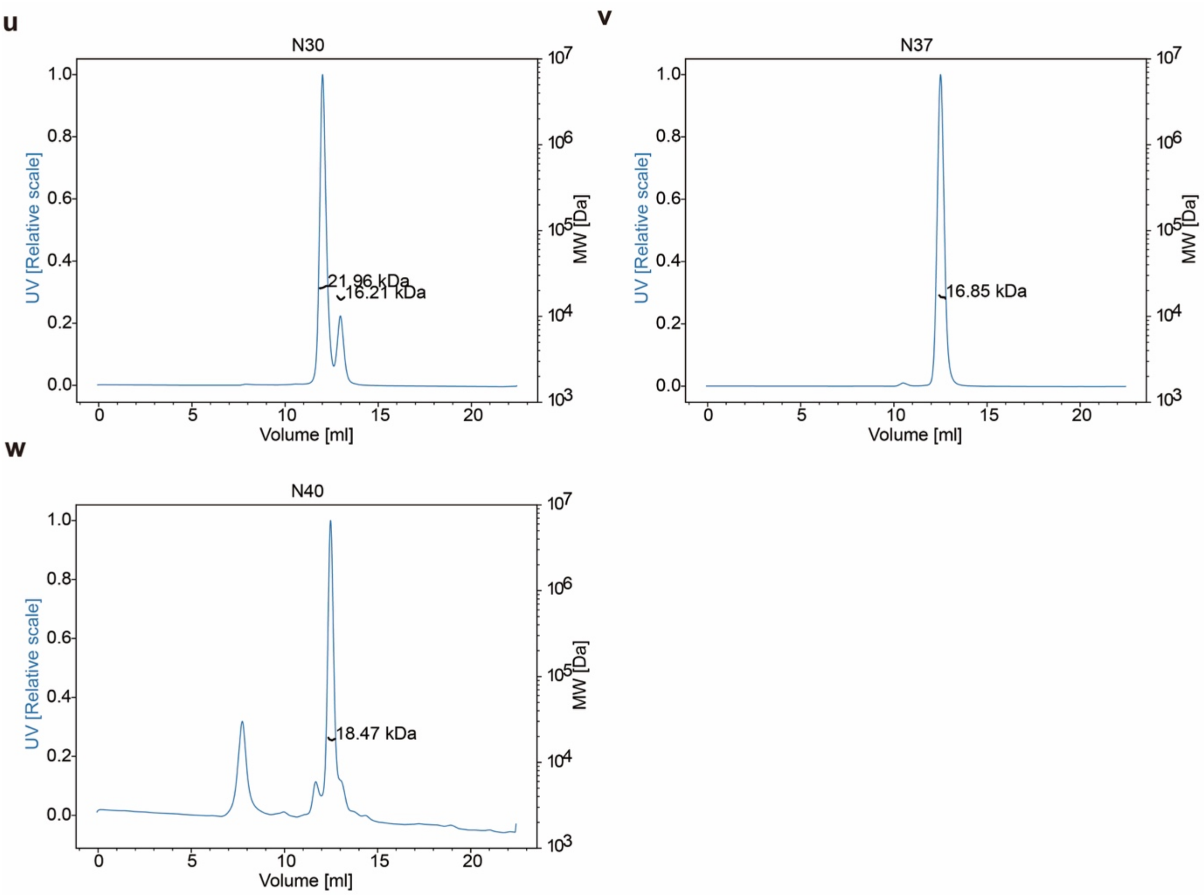


**Supplementary Figure 5. SEC-MALS of all dark fold designs.** All designs show the designs are oligomeric state.

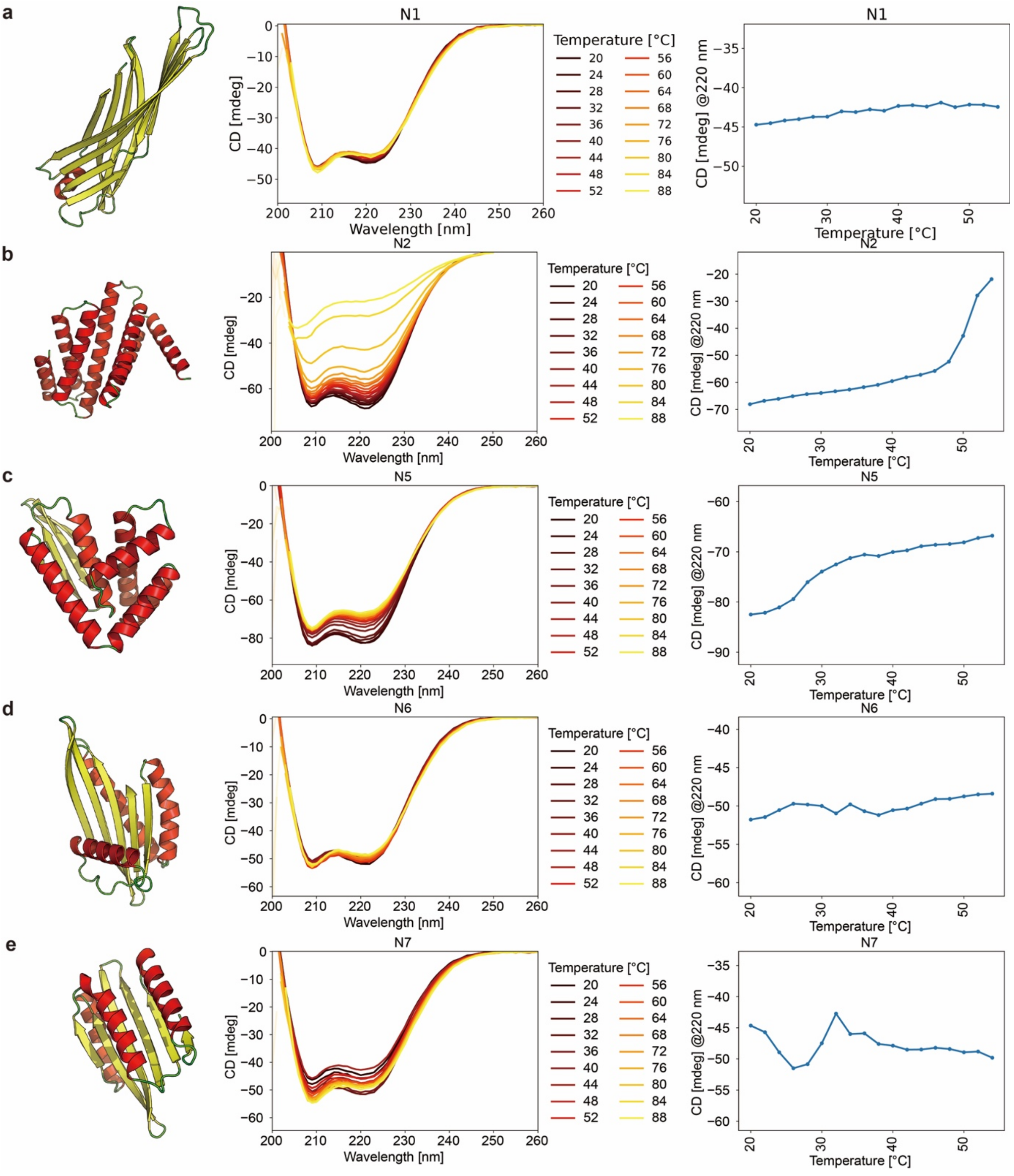


**Supplementary Figure 6. Design model, CD spectra and melting temperature curves.** a. N1, b. N2, c. N5, d. N6, e. N7.

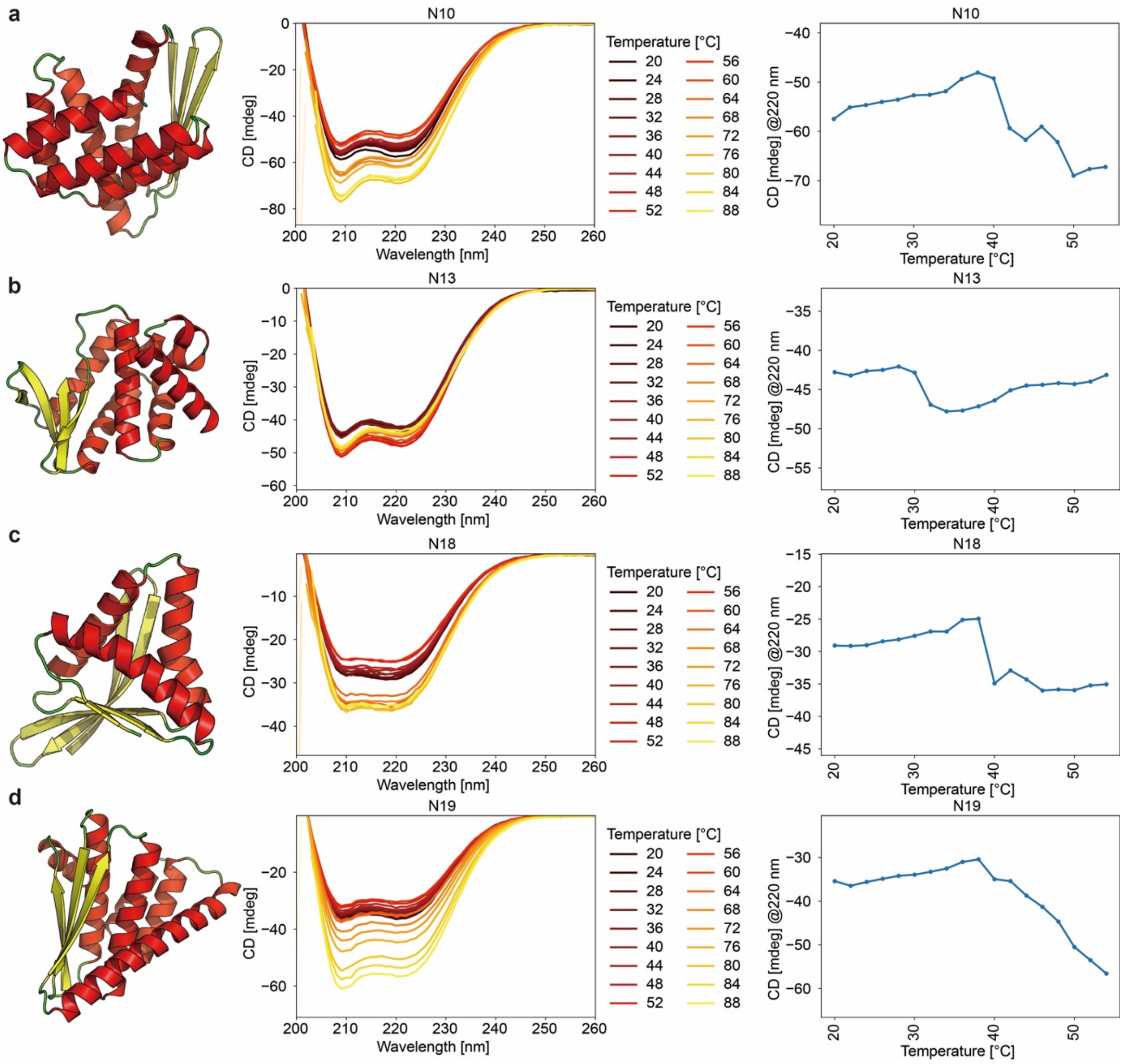


**Supplementary Figure 7. Design model, CD spectra and melting temperature curves.** a. N10, b. N13, c. N18, d. N19.

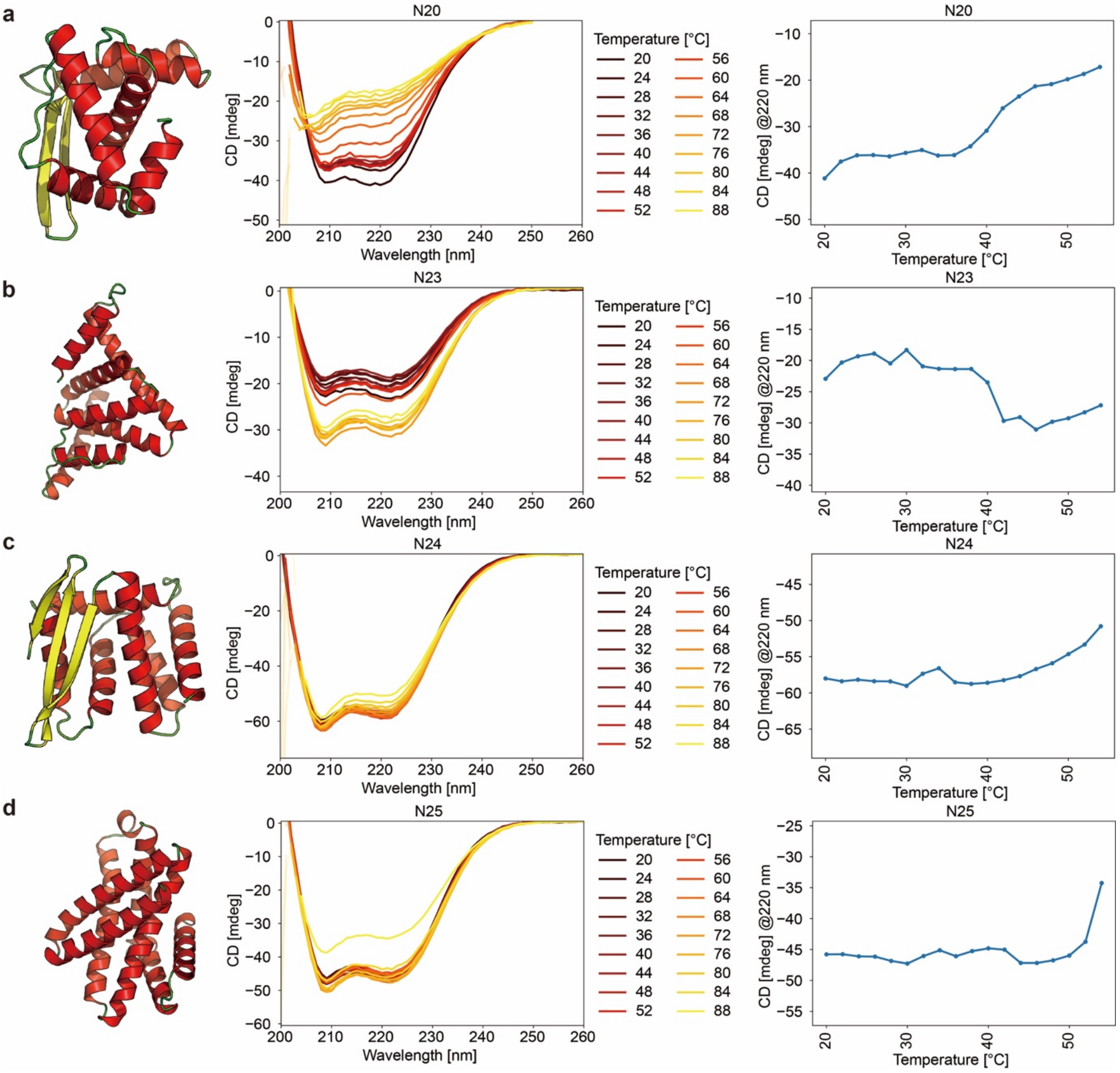


**Supplementary Figure 8. Design model, CD spectra and melting temperature curves.** a. N10, b. N13, c. N18, d. N19.

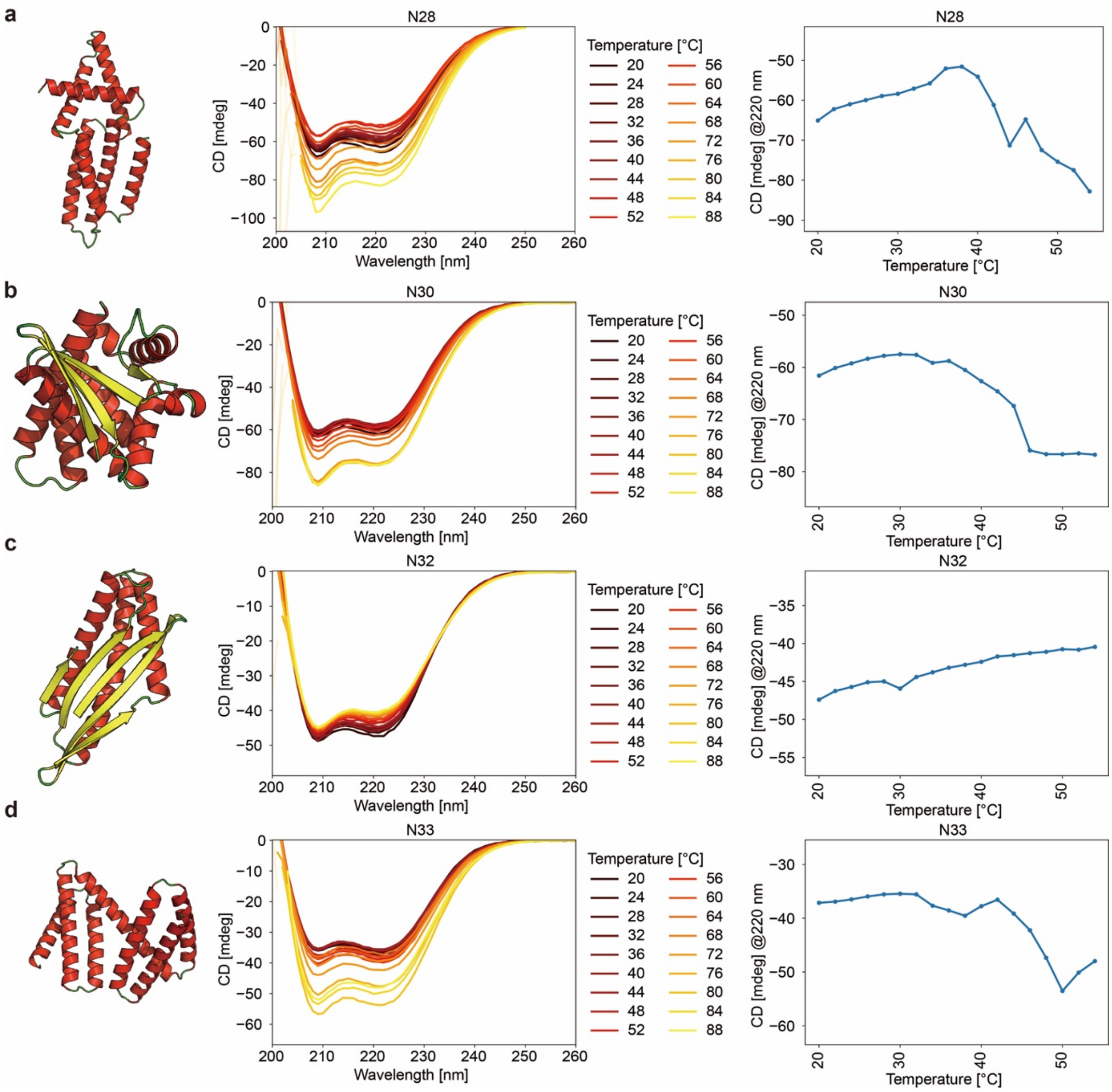


**Supplementary Figure 9. Design model, CD spectra and melting temperature curves.** a. N28, b. N30, c. N32, d. N33.

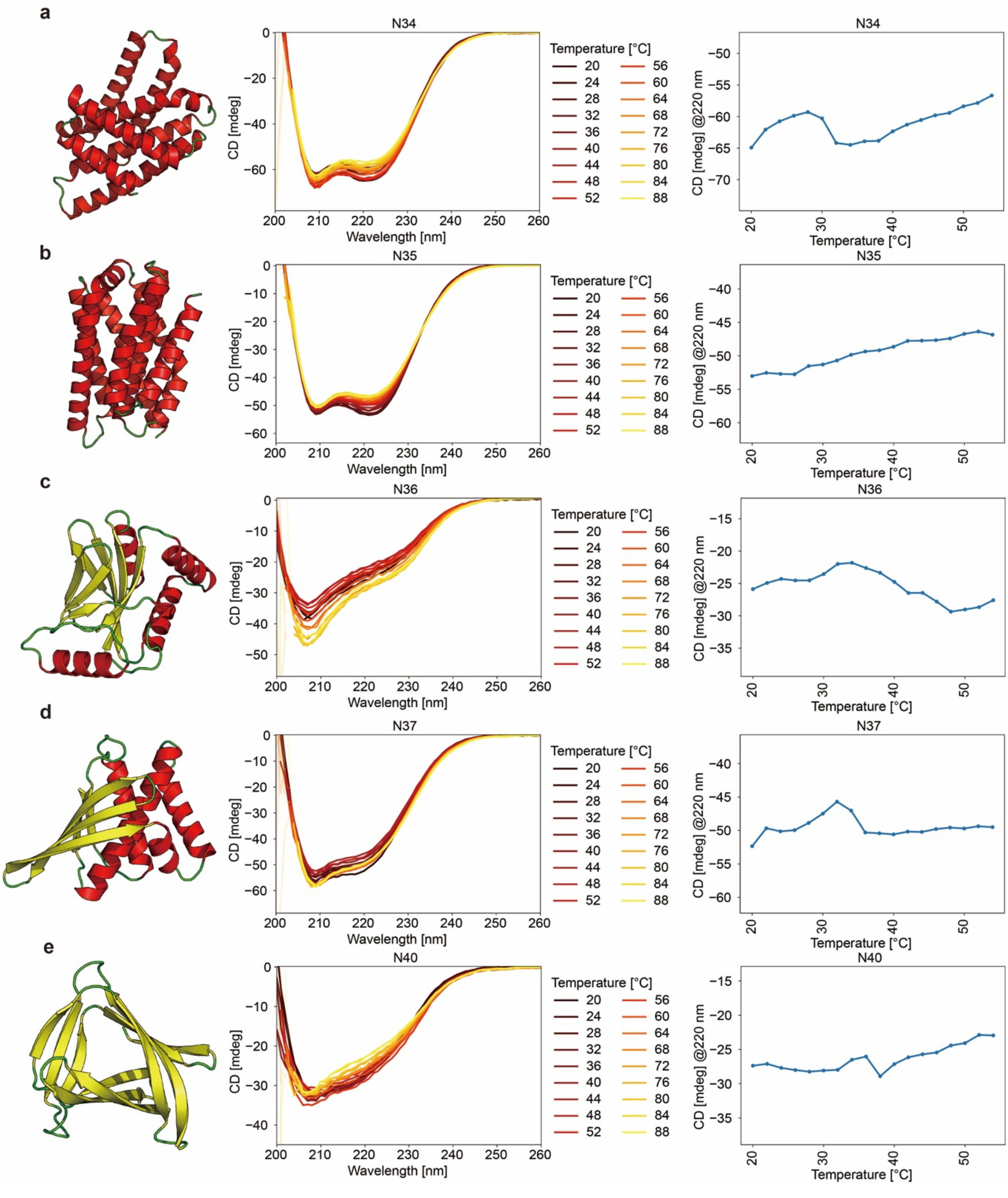


**Supplementary Figure 10. Design model, CD spectra and melting temperature curves**. a. N34, b. N35, c. N36, d. N37, e.N40.

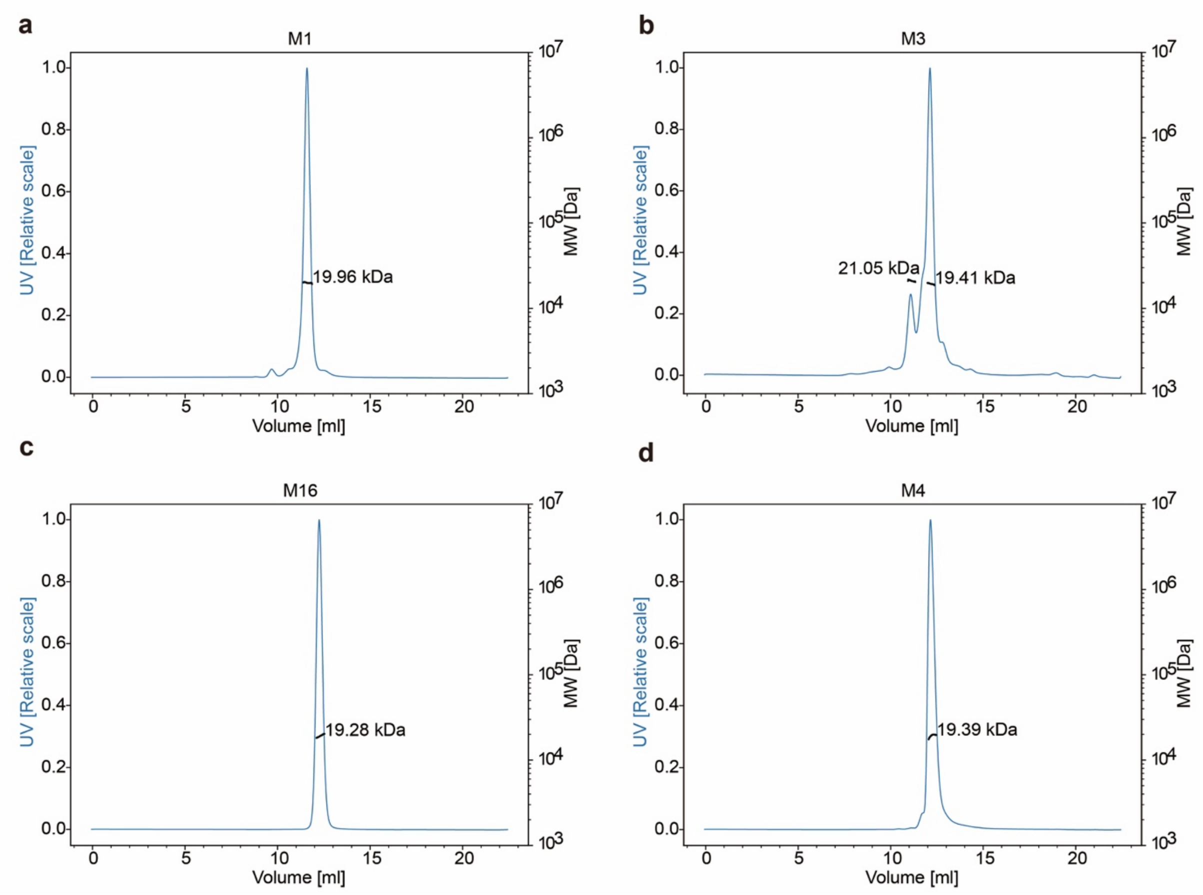

a
M1
UV [Relative scale]
MW [Da]
19.96 kDa
Volume [ml]
b
M3
21.05 kDa
19.41 kDa
c
M16
19.28 kDa
d
M4
19.39 kDa

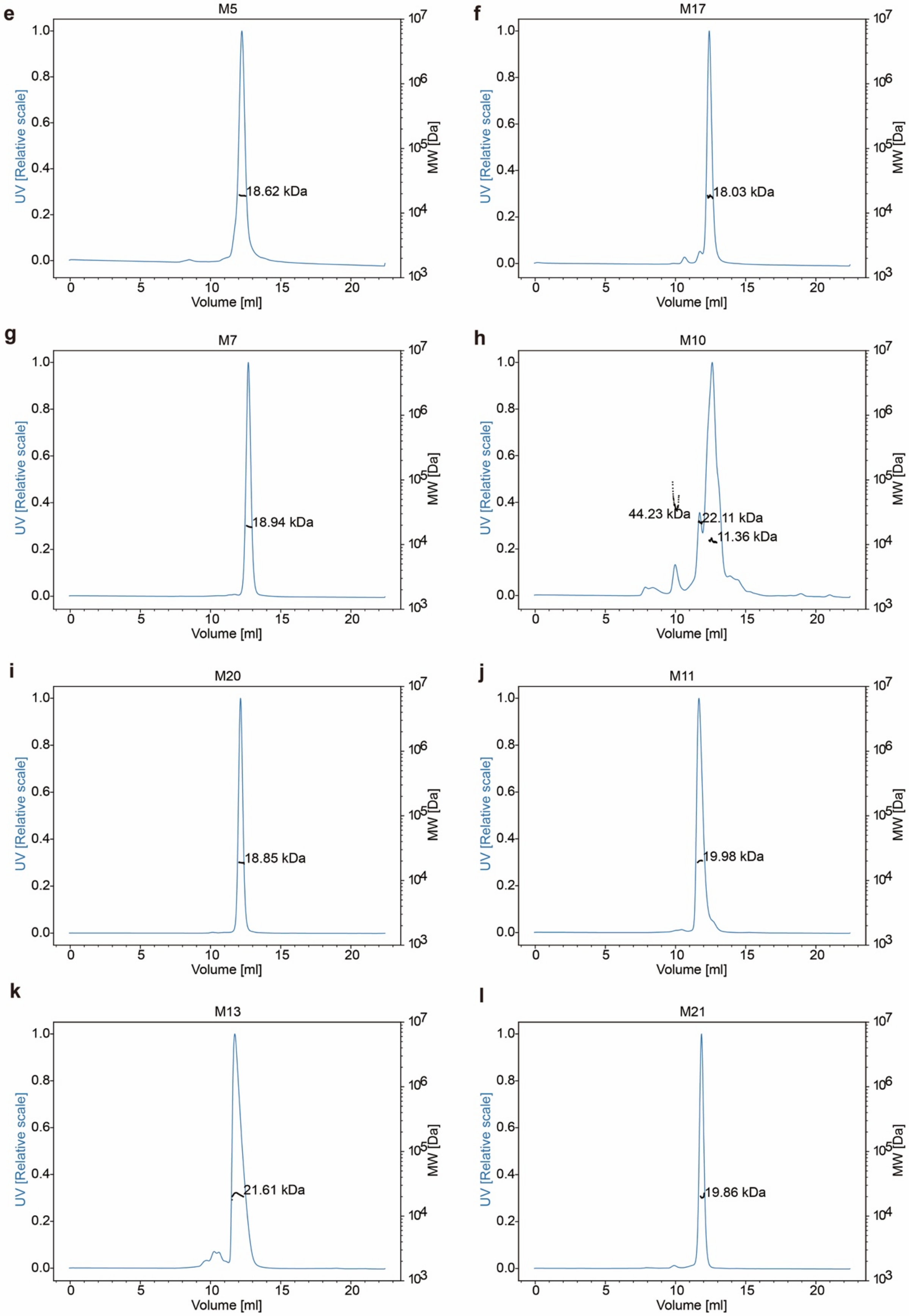

e
M5
18.62 kDa
f
M17
18.03 kDa
g
M7
18.94 kDa
h
M10
44.23 kDa
22.11 kDa
11.36 kDa
i
M20
18.85 kDa
j
M11
19.98 kDa
k
M13
21.61 kDa
l
M21
19.86 kDa
UV [Relative scale]
MW [Da]
Volume [ml]

***Supplementary Figure 11. Oligomeric state of the mirror fold designs detrmined by SEC-MALS.***

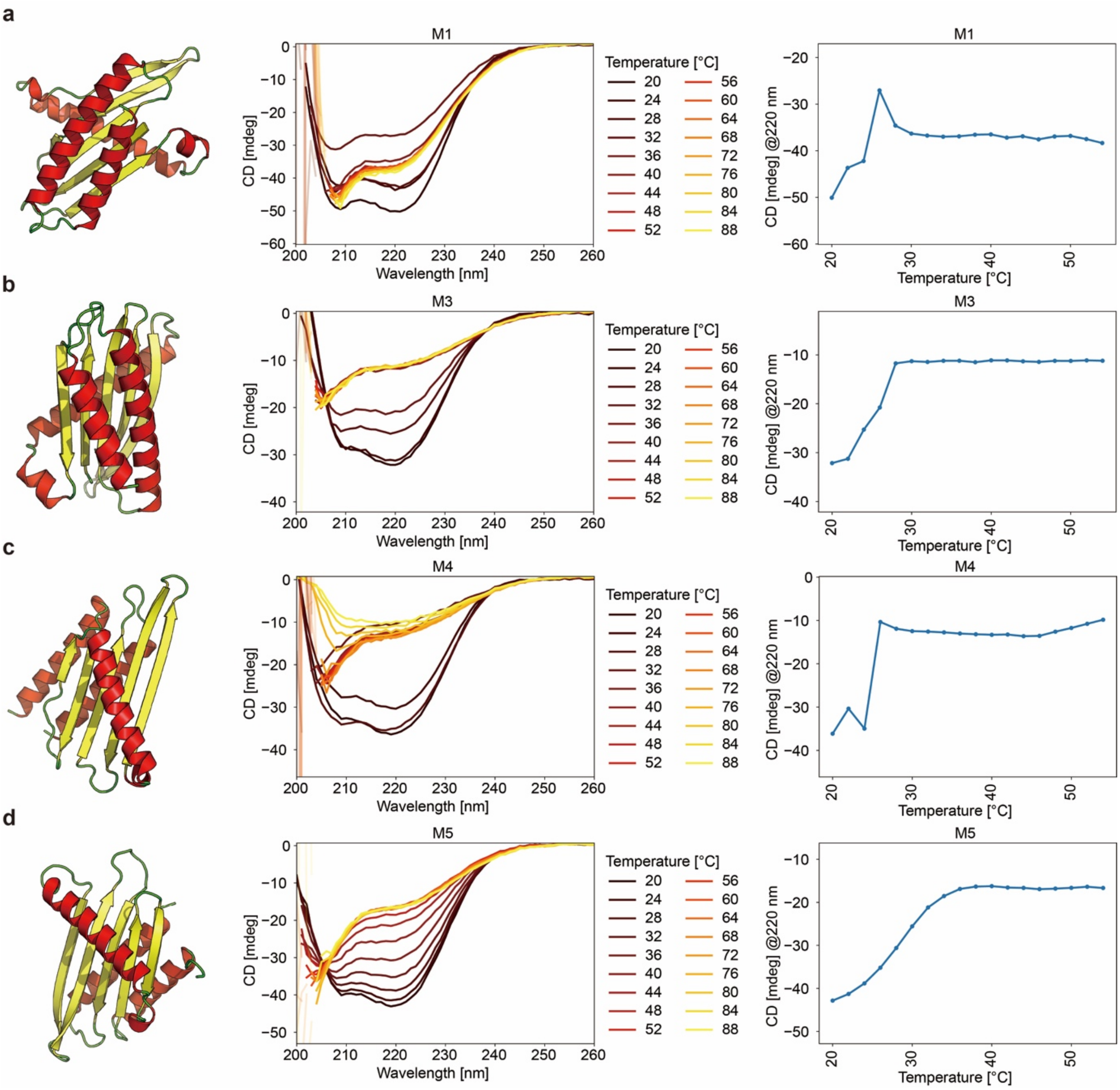


**Supplementary Figure 12. Design model, CD spectra and melting temperature curves**. a. M1, b. M3, c. M4, d. M5.

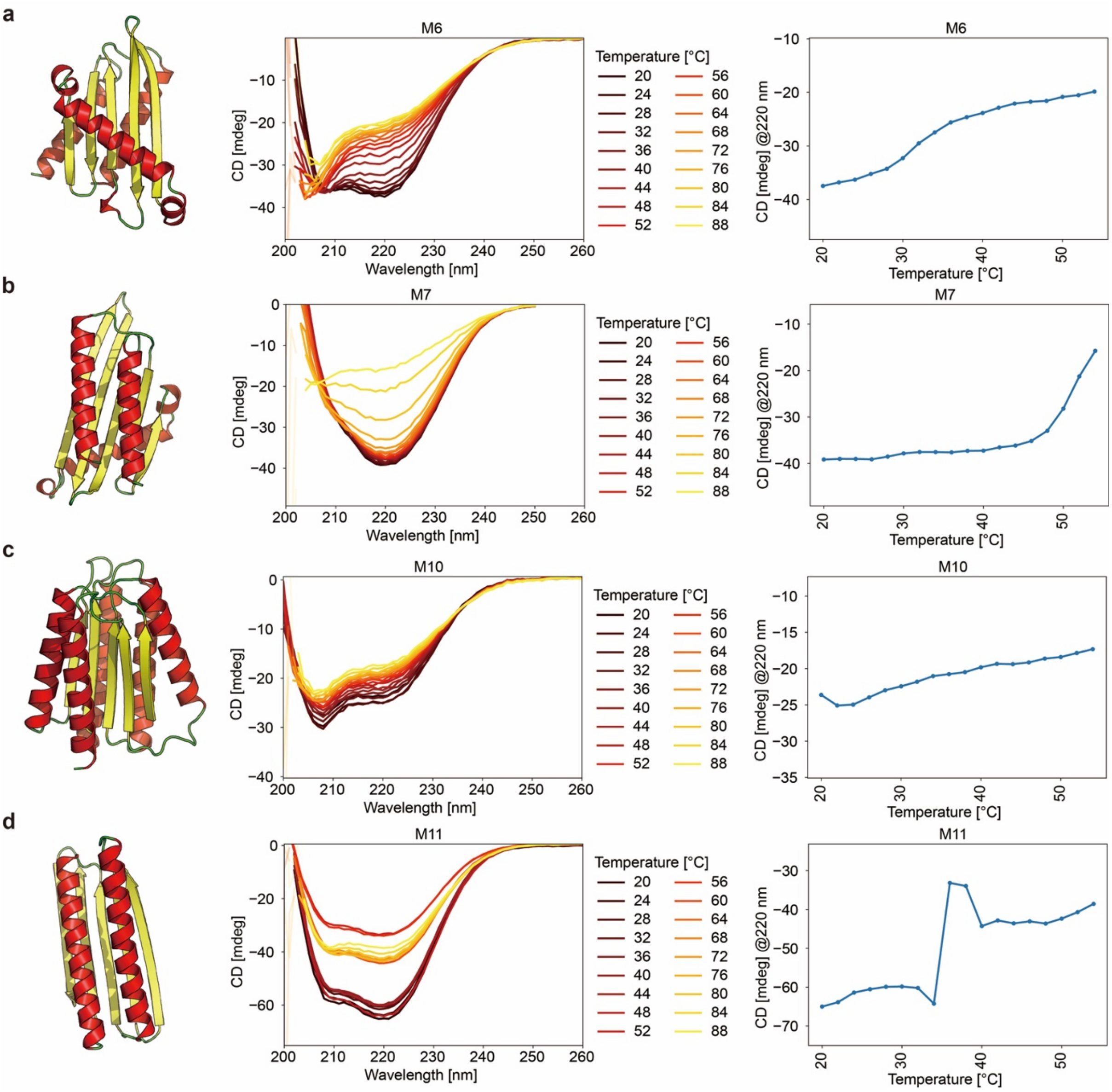


**Supplementary Figure 13. Design model, CD spectra and melting temperature curves.** a. M6, b. M7, c. M10, d. M11.

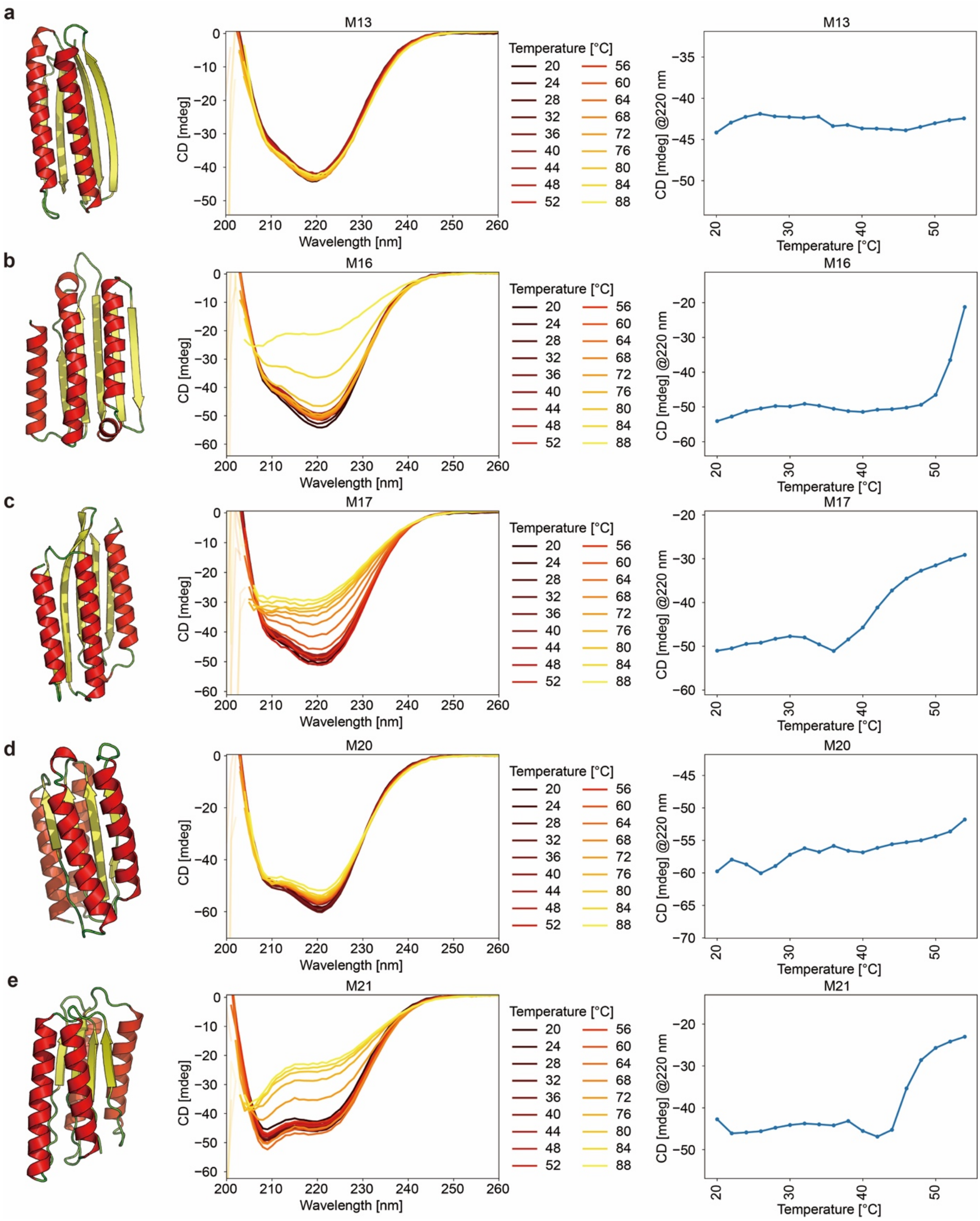


**Supplementary Figure 14. Design model, CD spectra and melting temperature curves.** a. M13, b. M16, c. M17, d. M20, e. M21.

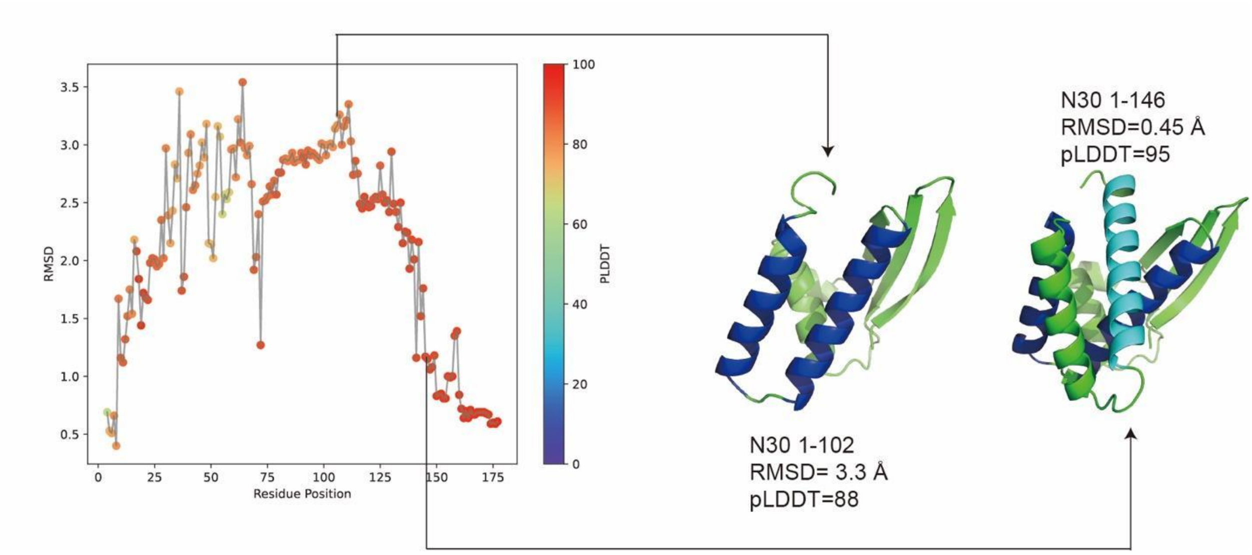


**Supplementary Figure 15, sequential folding analysis for N30**. Left : RMSD plot of Alphafold2 sequential folding prediction and crystal structure. Middle: prediction of 1-102. Right: prediction of 1-146. helix1 and 2 are blue and helix 3 is in cyan.

### Sequential AlphaFold predictions reveal a nonlinear folding pathway

When we obtained the crystal structure of protein N30, we observed a folding topology that appeared unnatural and potentially unstable during the folding process. This complicated arrangement prompted us to investigate how such a complex structure could assemble during protein folding. The most striking feature was the presence of helix 120-143, which appeared to be threaded through a cavity formed by two other helices (61-76 and 81-95), suggesting a non-trivial folding pathway.

To investigate the folding pathway of N30, we employed an incremental structure prediction approach. We generated a series of AlphaFold predictions for progressively longer segments of the protein sequence, starting from residues 1-4 and incrementally extending to the full-length protein (1-177). RMSD values were calculated by comparing each predicted structure to the crystal structure

Our analysis revealed distinct phases in the predicted folding pathway (Supplementary Figure 44 left). Initial predictions for segments 4-30 showed high RMSD values and low pLDDT scores, indicating that secondary structure elements had not yet stabilized. As the chain length increased, we observed the formation of the helix 1 (residues 61-76), followed by the helix 2 (residues 81-95). Interestingly, when helices 61-76 and 81-95 were modeled without the presence of residues 120-143, they consistently packed more closely together(Supplementary Figure 44 middle)., eliminating the cavity observed in the crystal structure. Only when helix 3(120-143) starts folding, helix 2 and helix 3 start to separate and form a cavity to accommodate the helix 3(Supplementary Figure 44 right).

This finding has important implications for protein structure prediction and design. Our results suggest that, at least for some proteins, it's possible to design more complex folding pathways involving threading and knot formation beyond the simple secondary structure stacking commonly observed. This expands the potential structural repertoire available for protein engineering and could enable the creation of novel protein folds with unique functions that rely on these complex topological features.

**Supplementary Table 1. Statistics of total generated backbones and their designability, diversity, novelty.**

| Type of fold | Total Number of Backbones | Designable Backbone Number | Cluster number | novel folds number of cluster represents |
|---|---|---|---|---|
| **7 helix(HHHHHHH)** | 182 | 130 | 48 | 18 |
| **8 strand(EEEEEEEE)** | 1100 | 666 | 117 | 10 |
| **low frequency alpha beta** | 6000 | 4265 | 2331 | 751 |
| **Mirror fold** | 24 | 18 | 8 | 8 |

**Supplementary Table 2. BLASTp search results.**

| Query ID | Length | Similarity Result | Query ID | Length | Similarity Result | Query ID | Length | Similarity Result |
|---|---|---|---|---|---|---|---|---|
| M1 | 156 | No significant similarity found | N2 | 177 | No significant similarity found | N16 | 103 | No significant similarity found |
| M2 | 156 | No significant similarity found | N35 | 207 | No significant similarity found | N1 | 172 | No significant similarity found |
| M3 | 156 | No significant similarity found | N28 | 189 | No significant similarity found | N4 | 110 | No significant similarity found |
| M4 | 144 | No significant similarity found | N39 | 216 | No significant similarity found | N3 | 105 | No significant similarity found |
| M5 | 144 | No significant similarity found | N25 | 185 | No significant similarity found | N27 | 116 | No significant similarity found |
| M6 | 144 | No significant similarity found | N34 | 199 | No significant similarity found | N5 | 157 | No significant similarity found |
| M7 | 144 | No significant similarity found | N14 | 183 | No significant similarity found | N9 | 160 | No significant similarity found |
| M8 | 204 | No significant similarity found | N23 | 167 | No significant similarity found | N36 | 141 | No significant similarity found |
| M9 | 193 | No significant similarity found | N12 | 231 | No significant similarity found | N30 | 177 | No significant similarity found |
| M10 | 193 | No significant similarity found | N8 | 140 | No significant similarity found | N20 | 139 | No significant similarity found |
| M11 | 133 | No significant similarity found | N33 | 187 | No significant similarity found | N21 | 142 | No significant similarity found |
| M12 | 133 | No significant similarity found | N11 | 184 | No significant similarity found | N29 | 183 | No significant similarity found |
| M13 | 133 | No significant similarity found | N37 | 134 | No significant similarity found | N31 | 144 | No significant similarity found |
| M14 | 151 | No significant | N6 | 146 | No significant | N15 | 171 | No significant |

| | | | | | | | | |
|---|---|---|---|---|---|---|---|---|
| | | similarity found | | | similarity found | | | similarity found |
| M15 | 151 | No significant similarity found | N26 | 127 | No significant similarity found | N19 | 168 | No significant similarity found |
| M16 | 151 | No significant similarity found | N32 | 146 | No significant similarity found | N24 | 159 | No significant similarity found |
| M17 | 151 | No significant similarity found | N7 | 124 | No significant similarity found | N17 | 162 | No significant similarity found |
| M18 | 152 | No significant similarity found | N18 | 120 | No significant similarity found | N10 | 169 | No significant similarity found |
| M19 | 152 | No significant similarity found | N38 | 135 | No significant similarity found | N13 | 149 | No significant similarity found |
| M20 | 152 | No significant similarity found | N40 | 108 | No significant similarity found | | | |

**Supplementary Table 3. HHpred search results.**

| Name | Evalue | Name | Evalue | Name | Evalue |
|---|---|---|---|---|---|
| **M1** | 25 | **N2** | 1.4 | **N16** | 81 |
| **M2** | 28 | **N35** | 1.1 | **N1** | 53 |
| **M3** | 65 | **N28** | 24 | **N4** | 130 |
| **M4** | 93 | **N39** | 4.2 | **N3** | 20 |
| **M5** | 27 | **N25** | 19 | **N27** | 4.2 |
| **M6** | 72 | **N34** | 19 | **N5** | 2 |
| **M7** | 27 | **N14** | 930 | **N9** | 43 |
| **M8** | 89 | **N23** | 15 | **N36** | 15 |
| **M9** | 170 | **N12** | 0.66 | **N30** | 22 |
| **M10** | 11 | **N8** | 26 | **N20** | 11 |
| **M11** | 1.2 | **N33** | 8.4 | **N21** | 25 |
| **M12** | 2.2 | **N11** | 52 | **N29** | 25 |
| **M13** | 14 | **N37** | 7.6 | **N31** | 1.4 |
| **M14** | 9.4 | **N6** | 4.8 | **N15** | 20 |
| **M15** | 54 | **N26** | 1.4 | **N19** | 4.2 |
| **M16** | 2.3 | **N32** | 1.2 | **N24** | 1 |
| **M17** | 72 | **N7** | 14 | **N17** | 6.1 |
| **M18** | 18 | **N18** | 6.7 | **N10** | 10 |
| **M19** | 490 | **N38** | 13 | **N13** | 20 |
| **M20** | 17 | **N40** | 17 | | |
| **M21** | 0.13 | **N22** | 55 | | |

**Supplementary Table 4. Information about all designs with experimental validation**

| Rename | seq | Scaffold name/ SSE descriptor | pLDDT | RMSD | Max TMscore |
|---|---|---|---|---|---|
| M1 | MYWIVEVEEDENGNIKKVYVKYWTDEELKEGKYSKEELEEWEKREEELIKETKKYEEENKEDESKKELIDDRKRDENGNEKRDIYIPAIKYENKGYVSYLRLTEDTDEEKRKEYLEKSKESLEKILDYLKSKKDKGVTEEDIKKVEKVLEKVKSKI | 2j3w | 0.849 | 1.600 | 0.445 |
| M2 | KSYRGSIEYDEDGNLNIHKPVLAETEEELKEKLTEEEQKEYEERRKKFSEEYEKAKKLKEQGDKSVYSYHYKLTDENGNTKGEIIYITDEKDKNKLKVYVELYNTENEEDKKKQKEALKKVIEKLTEWYNKYISKNKDEETQKKFEEFLKQLESLL | 2j3w | 0.877 | 1.630 | 0.444 |
| M3 | MYVYHRLEVDENGKVLYYDSKFYKDEKEYKENISEEEEKEYKDYFEELYNKSKKLKEEKDKNYTIHYNEKEYTTKKDGTKEISYGLVHIEHKDGSLESAIIINPSKNKEEEKKKLKKAKEHLDKVSEYLEKRLDKGVPQSTLNSVKELRDTISKYI | 2j3w | 0.928 | 0.852 | 0.452 |
| M4 | SKEEKIKEFKEKIEKTKKEIKEKKKDYYLEYEVKDGEAKEPELKEISDEEAEKKIEEYKKYRDELKKSGKEKDYNIKEEEEEEYNEETKDTSRYVYIHIENKKDKKNRKSLVYYKSNKKTKNEELFKKFEDELEKIYETYKKNL | 1sko | 0.899 | 2.131 | 0.392 |
| M5 | KEKKTDEEIRNDLKKAKKNIDKEDISLRLTIDKNGEKKYSYNDESKITEEEKKYYEQTRKKLKEDYEKNKNSDSNYSYYKKERKNSKTGETNSLEILKKKSSDGSSKTEYYLYLKISSSDEDEEFKKEAKEALDESIDEVDNSK | 1sko | 0.886 | 2.339 | 0.395 |
| M6 | SLEEKKKEIEELLKRIKEEYESKAFTAYLYYESNGKMEYATTEEELNKEETQKKIKEYKKKAEYYENTEELKKKGVEVEKEEIKRTENGEERTDEYIYIKDKENPENNFTLLTINRKEPITDKEDKEKLKEMIDKYEENLKKQL | 1sko | 0.915 | 0.703 | 0.383 |
| M7 | SEQEFLDKAISYLDEARENLTKRPFYYYHSMDEEGNSKSANSEEEYKKKETQDFKERVLKKLEELKNSKDKDKYEIKEEKYEYTEEENGKKKKYKIEYRYARKDGKLKAYYLEIYSSDKMKDEKTIKETKERLDEIKKELESLR | 1sko | 0.944 | 1.082 | 0.373 |
| M8 | SLEEVKNAQKELADYLKSDSNSENFSYEEEMTIKNTQGKENKYSRKLEVNKETGEVKTEYNLEWQGSQEQKDNLLQLEKKELEELKKIYEKSSSTDEIITETISMDSSGNVTSDDESVKKSIQDLYNTYKSKLDQGYDINQKEEKLNLSSGKDKSLDYSQSEVSSTNKNLTININIRLPKSAYEHAKNFIINHFQAHKSFLENS | 1914 | 0.918 | 0.800 | 0.379 |
| M9 | MIVYLNLLYSTSREGHEEALEMIKETIEELKKGEKNIKKIKLNLEDVDEELKKKTKEKIEESTKKIEEKVEKNNVEKYVY | 2lci | 0.914 | 0.769 | 0.406 |

| | | | | | |
|---|---|---|---|---|---|
| | NIYLSADSSDRESIEEYLKFLEKVTEYILKEIEKGNSVSVTYINFTDSTETEEEKRKKTKEKSEELLKKIEAQSETGIDEVTVLYTPSDEELSNETKKLVEELKEEIRKEREK | | | | |
| M10 | RERRIETDGEVNEEDTKKAIEKLEEANKKLKEGDKYVSYSEISHTSESKEYVEKIRKEWEELNELSKEYYETTGATVSNILYIDNLKKEHKEESEKLTELIKQWRKLLQKAKEKGTKTISFHLVLGDKLESRKKTLDASEKWIKELEEDDEEWDGDELSYYSISKPSKNEELRKKYEEKQEELERLLRESGES | 2lci | 0.818 | 1.460 | 0.396 |
| M11 | KNSLEINVNYSYENGKETEIKTEYKLDLPDYDEEKKKEIKEKLEKKLKELKELIEEARKKGIDGLKINLNFKSENVNEEDFDKFYEKLFKKIEETKEKIKEALEKKKPLTLNTNLEYKTEKDKTIYKEEIDEK | 1qys | 0.842 | 1.782 | 0.508 |
| M12 | MFNLKAKLEYDGSKGEEPKLEIEKEIDLNEENKEKLEKETKELEESLKKLKEILEEEKKEGRNDIKVKIEISITNATEDEIDYVIKKLKEITEKIKKLKELLEKRKNGSSEKYEYEEEEEKNNTKIKTKIELK | 1qys | 0.889 | 0.874 | 0.499 |
| M13 | SVKLDIKSDLHSENGKITKDESEYKYEVDEEYKDELGKIVEELKKNVEELKKEVKKLIEEGRKNVKLEFNLKIENTKDEDIEWIKEKLKEINKEVKEITEKLKKLSSNESYKYEKKYEDENIGFKYELKLEVE | 1qys | 0.927 | 0.623 | 0.426 |
| M14 | SEEKEKIEEIKKETKKKIDSGDKNVKTEEQEIELDETTTDEEVVDLVNYIYDSIWENEEYISKKKKKNVSLEYKYKITKTRKDGSKTIIEIEYKSYFKDTYTFSEEEKEKTRERRETMVRTSLETKKDQEEEEEYKSDNYEYKEKLKITHE | 4gt8 | 0.943 | 0.631 | 0.500 |
| M15 | SELLKETEEAYEKAKKKIEENPEHADITINFNKSGESISLEDIEKSIFKLLDLSSYAVKSKNEDKIKKVNLNINLNVNTTYSDGSTLNVNLNITIENPGDYEEWKKFLDKKKEEIKKTFDEIKETKKPKTYSYSYSDSTTKTSLNLTINYK | 4gt8 | 0.895 | 1.315 | 0.467 |
| M16 | SESKKLIEEAKKEFDKQLAEGKKEIDIEIEYPKKKYDENYKEALEFIKLNSELRVEYVKKSREAGVEKLTIKTKVKLEPTENSDHKLDYEYEEEMSKKALEKTDEKEYKEKLEKEYKKIEETVESNKDYEYEEEYEDKETGYKYKKKVEIK | 4gt8 | 0.933 | 0.755 | 0.469 |
| M17 | SKEMEEFNKEIEEYKEELKKSKEEINIKFPYKKYTENVSYEELKTEKTKLIDLLKEIYNTTKEKKLKKVKFEYKREYSYKTEDGSEVYIKLEISASNLNKETNYEEKTNDFEKKFKSSLDKAEKSTEDLSEEYEYKDDKSGLKLNYKLERK | 4gt8 | 0.901 | 1.111 | 0.428 |
| M18 | KRTHSLNLDIDIDFKGKDKNINKEEQNEIKETLEEQKNYLKTLLENADTGKTVSYNSYKKEYKSGLTINKSGDDISSDSTETKTLSLNINYDDKKTGNKLNINLSLKLTSIKDSRETIEKALENVEKIEKWIEESLSTGKSYKYSYSYTEEK | 2iii | 0.908 | 1.674 | 0.412 |
| M19 | SNKKTIKSEVDITRKKTDPKRDKEMKEKTQQKLDQNESNTNNYLNNIDNYD | 2iii | 0.881 | 1.977 | 0.502 |

| | | | | | |
|---|---|---|---|---|---|
| | KESETENKTEETEDFTINNNTTYTKE KDNKKYTYNQKTTITDKKSGSKVNINI TITNTDKTWSDEFKQKYNNSYQNIKN SINTSFNTGKNIKKNNEESLEY | | | | |
| M20 | MKYYESIIKIGVSEEEQKELLEKTKER IEELRKKIEEGSDEEYSYSEISYDESL SEEHKKILERIKETNKKIKEKLKEGDV YLSIYITSSTELVKKEDLEEILEKNEEY LKKLEEGEEKPYYSYTKLKGVSKELE ESAKELEETVREVYNKT | 1abe | 0.844 | 0.875 | 0.450 |
| M21 | KIRSTLLIILKASEETLKETIEIIEEELER LKELREQGRRERTSTVKLTLSDDVP EEDREWLEETIREIKEELDKLSDDDIP GPIIVTYSNDGVSEEEKEEGKRTLEE YREELRERKENIPSRILLSNLESEELE KELREKYEELDQLLRSL | 1abe | 0.874 | 0.735 | 0.430 |
| N2 | SKKELEEREREKRKELAKSKTEESKK KLLDYYVESELQIEEKKEKGEPESEV EEEKEKLDKEIDEDIDIYYEDKTEEEK KKEKYNFKTEIYREVLRQRALRYASK EELTEEKIRELVRELKKKMDEIRDKY KVKSKEEVDNEVRRMSYETADYLHD EKGLLTEEQRELWRKVVEKEV | HHHHHHH | 0.916 | 1.569 | 0.482 |
| N35 | EEEEIEEKLEKYFEIVKKYEEEKVKAY QESKSVEEYLEKYYEIDTKRFKELIDL YLSDEEDYEEKSEELTKIYNETREKY RELDEKYRPYIEKWLKTMTREQVEE YAKEVLKRLEERREEIREKLKELKDN REKLMYLYYEYHRVDYRVWVKRLVE KLGVSKEEALEIYEEYIEILEESLVKS GVITEEEAKELKKKELEELKEELK | HHHHHHH | 0.944 | 0.634 | 0.477 |
| N28 | SEEELQKRIDEKKKKYKDYVESGKPT EWYKKTIELLSLYRKLKEAKKTLKEKL EKGELDENVTEEEVEELIEELEETLE ELEKYFKIKGISDYDSLEKQYKKNYE KIKEKEEAESKNDESKKEEVKKFEEE LEKQVKYEFEISKKTSAEDRLLEKSK KVLEKAEEYLKNDEKYKKYKEEVEKE LEEKIK | HHHHHHH | 0.906 | 1.095 | 0.482 |
| N39 | EEEETSLKLYRYIIDTEIEETIERGKKS KKKLEKEGKTELIEYVKDRTRELIKKH FKDDRSEEEREKEKEEAEKESKKRR EETKKYLEENPEYRKKVEEWLERRK KIDDEYYEELDTLPEELRNSLKYSDS LIAERLRIELYRLEKQGATLEDYLKAY TKSLELQKEVHSEILKYYREEKAEEL GVTKEELEEKEKKLKEIIDKQEKLSKE IYEEL | HHHHHHH | 0.944 | 0.703 | 0.454 |
| N25 | SEELKKLIEKTEEELRKTTDEDEEKLK KASELTRKFYFSEEDGKGKENQKNIE DSTKYLELKATGTIDTKTTEEEKKKLY DSSVDLQYNLMKEIIELNKKYSEEEK EIVDLYSLLSVKTYSEYHYNKLLKYVD KEKATTLTYEYSKESLKKASEILKNEY DKETVEEITKKINSYYEEWYKKRTS | HHHHHHH | 0.917 | 1.474 | 0.474 |
| N34 | DEEYRKKLEEIFQETTKKVYELEKKK DNLSEEEYEKERQKIRDESKKKISEL AKEKGWEEEEIEKHQKYYDLEYNNL WKTIEDYYEKISENKENEEKILENLSK YHYYLYEALLLLSKTKNLSEKEKEET | HHHHHHH | 0.935 | 0.954 | 0.427 |

| | | | | | |
|---|---|---|---|---|---|
| | EKEINELLKKTKESLKKFYELLGLDFEKNYEKHLNQYKKISEEEYKEKKTITEEDLENTNKILEKLKEI | | | | |
| N14 | EEKEKEIKEYEEEYKKELEEKGETDKKTIEKKKEYTRELLEYKYKHKDKSYYEIYEELSEEKTEELVKKGKSYETLKEELEKLKESSLYIIKKEKNLNEEEKLKVKSKLDEIETKAKEKYLKKKGKSYEEITKLKKEKLEKEEEEKLKYYKDEKKKEEIKKEYEKKKEELDKEEKKKKEEEKK | HHHHHHH | 0.916 | 1.760 | 0.468 |
| N23 | SEEEEHKEEIKRRSEELEKELKKYEEEGKDPIRSYTRDELEKATGEEVREDITPEEALERLIEEMAKRLGLDEREKKKVLLGLYEEIDEDSKYTSRETATLIQVRKLSRLLVEYKKDPERTKKEIKEYVENRIKRAEESFSKKFNGDKTVTEEEKEKWLEEELEKHK | HHHHHHH | 0.935 | 1.173 | 0.450 |
| N12 | SREEIKEKHEKKLEEELEELDKKYKEEYGVDEETLKEIREKTKENWKLLLELLDKYAPEKYYEYFLETITKVLILQYKIQLQRYILETKYGVDKEEFNKELKKEYEEARKEAEEYLEKLKKSKDKSDEEIYEELKSKVEELLKEAEKLTEEAEKLLEELEKGGDEETKEKAKEKLIKASRLSFLIIYLTLYSSAYFKARETKKTEEEKEKYLEEMKKEIEKTMKKIEERSL | HHHHHHH | 0.955 | 0.622 | 0.443 |
| N8 | SEVDVLLLILDELVKKSKDKEYAEKKWKETKEELKKLKTEEEKKKLLKKTIEELLKHEGKKEEEKEDVKYLSEKLTETYEKLEKNKTEEEKKKIWEETEKELKNYSKYSYELYKEIKKRVLENTGKLTGYKEYAKKLLEE | HHHHHHH | 0.943 | 1.016 | 0.495 |
| N33 | DKEKEIKKLYKEYFETEEKAYQAYKNKDWEEYYKSYNKLIEIKKKIEEYKKSGDSEYKKSIEKAEKEREEKKKKLPEEVRKFYDELEEKINEELELEKSLEEIRKKSESGSELDKLRKETLDTYDEVSKLKEERAEKEKSSPEEKWKIYEENIKKKKEIIESEEDEEKLKKLLEKLKKEKEKLKKEE | HHHHHHH | 0.915 | 1.549 | 0.449 |
| N11 | SKEIEQLKKEYEDKYNEWKNSGFLDKSDEEQIKELSDYLEEKIKKYTGIEELSEKTKNEIKESVEELYREHKKKYDDKEEQLRTTLEDILITVLVLKRLKEEYPEITNEEITYSLKVITRLRLWDIRNEGVSEEERRKKREEVIEEVYKEYLEEYPNISEEEKERFRKHLEKTAELYEEVRKEL | HHHHHHH | 0.954 | 0.802 | 0.487 |
| N37 | SVEVKVTQIDDETWKFKTTITKENGEKEEKESTITKEEVKESYESEEEYESTKERIRKKFEDLSEKEKYTLSNLVLAITKKLKELYEEYGAKEVKVTIEPINGKPLDKETKERIKESIEELLKEKGYDVKVELE | EEEHHEHE | 0.940 | 0.660 | 0.489 |
| N6 | SKELEKEEKSEEDGKTYYKLTVTKVSLRERNGKKYLFISEAEKTTTEETKEEDRKELKNITEEIMSKIEELWEKGLSVEEIKEKLKKELTPKYSEKGYKVEVELLHKEELDDEENRKKWEEYKKLAKEAGIEDVDDVETILHVKIK | EEEHHEHE | 0.937 | 0.939 | 0.403 |

| | | | | | |
|---|---|---|---|---|---|
| N26 | STEVILLRDSRTGLYVLVERETKEEN GVTHVTITVEVLKDEETARERARRLA EERGAKLTELPEQDPVDYLYETVKK NLEEGNKVTVIFRSDPETSDREETER LYEELSKRLKEEGYSDKVELRYEE | EEEHHEHE | 0.939 | 0.701 | 0.403 |
| N32 | MKFKIKVSVNEDGKVSVDLTLEETRT ENGEEKKKTINLNEKEKKEKLEELYQ KALDNLKKGTEEDEKKAEENIQKAM DIIYEIIDKKIEEYKKKGYKVKLTISYEL DLPEDLTEEQKERLKEWTTKNLQKV KEKLEPKVESIELET | EEEHHEHE | 0.931 | 0.588 | 0.470 |
| N7 | MELKVDINSDEDGSKLAIFTEYKYEN GKLKKKVKIYYNNEKAKKKAEELKEE YKNKGIEVELEKVEEEKFYDKVKNYI EEYLEKNKDLKKVEFNINEPDEKRKK EYEEYIEELKEKYKNIEFTL | EEEHHEHE | 0.935 | 0.548 | 0.449 |
| N18 | TTKEEEIEIEYKGVKYSEKVTFTTDEN GTTDGEINYNSEEEYENTYKNFKNK MKEKGLSEEDWIEALTKLFEEIFKKA NGKITIKINLKDKKKQKEYYEEISKDV KKVEKKLGREINVET | EEEHHEHE | 0.944 | 0.828 | 0.467 |
| N38 | KKLKITISEDENGNLKLTYTEITKDEET GKETEETSEYKDENGNYKLDKLFDE YDEWSDLPEEERKKKLKEKLLKYIED VIKGTNKEYDEIYIEVEIKNGSNKERL ELTNELLEEEAKKKGWSIEKNGNKTT YSN | EEEHHEHE | 0.933 | 0.774 | 0.424 |
| N40 | KEYTYTYTTSDGSTVTVTIRINDDGTI EIYTNREPDTRRIPDRTRTITTYDKET GETTTMTITIRSKDGGDITERRTRSE EDSTWSRVSYEDATLTITLSDGTTYT EE | EEEEEEEE | 0.919 | 2.076 | 0.467 |
| N22 | EKKEEKEEERKDGKKIKKKKITKELN ADKVKVTLTITITDKKTKETEEYTYEL NENKTYEKKGNGKTYTYNVTVTDEK TKETKKTTITITTKSSSEEDTEIKEELK IEGEGNEKNTVNIEIE | EEEEEEEE | 0.901 | 1.183 | 0.474 |
| N16 | SISVTSTSSTTVTLQLTDDTTKETTTT TLTVGERTEVETKVTTSTDSNGKTTT TYSFKLAGHTVTLTNGSKTLTSTNNG TTHTATLTIEKDEETGKYYLKVSN | EEEEEEEE | 0.904 | 2.648 | 0.477 |
| N1 | SSITVTINVTINFSKDKDGKYRVTVNI NITVTETWDDGSTDTKTANYTDSAE VKQSKTDKNIYEANLTGPDITLTLTKT TPNTTNTSDTTTTTTTTSTTLTKTYED KEKKRKITITTNITITTTTTTTDTTTTTT TTVTIKGIKINIKQSKKTLEEDIEKKTT YNYTITYTK | EEEEEEEE | 0.915 | 2.570 | 0.450 |
| N4 | TETVTDDTTGETGTATLSNGKITGTG DLTTKTVTRQELIESKETLKLEETTTK TENGETITETKKQEITPEDLKNYPEE WTFYTLRVGNKTKTWLEITDENGKT HYYKLL | EEEEEEEE | 0.906 | 1.441 | 0.427 |
| N3 | MKEETKDEESEGKKTRTTTVQDEKD GSTVTLTETSPSTVEWKDLSIEVERN SDGTITLTLTGGDLSYSSDGTITRRD SDTTTTVTITVKEGQTLTWSDGSTTT VQ | EEEEEEEE | 0.904 | 1.025 | 0.468 |
| N27 | STLTINVEINLTDDNNNKYTASGTATL TSLGETETMTITLTDSNGNTKTYSYT | EEEEEEEE | 0.926 | 2.474 | 0.487 |

| | | | | | |
|---|---|---|---|---|---|
| | FTKEDVDSKTINIKLEKSGLKKGDTVTQTQTVETTDSNGNTITKTYTLTVTYDENGNINISLS | | | | |
| N5 | SREETRDDDGTRELRYKNPFDGSEITIRSPRSEDFERVSTEVLLLELLETIGLLLSREELEELIKLLQEEYPEWTPDQALQWLRDTTEKTIKEAAEQGWNKEQAFELLLSSVPEKIAEYLKKLKEERETDEERRKQTEEDREEWEEELRERGYTISR | EEEHHHHHE | 0.931 | 0.926 | 0.467 |
| N9 | KYKYTSKSSKGETIEIIIYKKDGEVHVDIYDGDKKITLEEAVTSYLLYLYRTHYKSYDKSDEEIWEELSKEYLKRKYSEEERKKKSLEEVKEEIKNEFLKKETYEEEIEYLKSLYDDPEKLKEYYEEKKERLENSALKPYYETAKKKTKEEGKELSKLEE | EEEHHHHHE | 0.935 | 0.737 | 0.411 |
| N36 | ISYEKLDDNHSALIYDSEGTVKELIDEFKEKLLEVLGSEEKYEEFWNYLKSDPNLKVVVGLKGFKLTYKDGKVYFTADKNNKEQQEELKKVKDAGGKVEGNTLTAWTLDEAPLDYEGNWRLVILGDKSRHDTLKELYEKLK | EEHHEEEHEEH | 0.933 | 0.858 | 0.416 |
| N30 | SLIISERKEEGETVTWDLSLSEDSNENEKKAWKRYFERYGLTDEEISKIESIRVEGTEEEVEKMYYYYKLELEIREKLNSEETEEKLEEIWRLSSKGTEENLKEAKEIIKELLKEIGYKEDVEKKAEEYLEGLQKYLDYLSKKFGITREQLGKRETRSKLYRESLENPEKYPLFKLK | EEHEHHHHH | 0.931 | 0.931 | 0.487 |
| N20 | SSYEELQEWLRDGNREKFKEVLDELEKQMKKTGEKLEISTDKETLYDALSYTYSHRSEYDDETRWKFDSLLRILAVSSKYRGNYTSEERLEYYYKEEGKSEEERQKEVDRYKEYAKSNKDGKVTLTLTNEKGQKTTITW | HHHHHHEE | 0.932 | 0.825 | 0.448 |
| N21 | SEEEEERRRRLERLRRTRPELVKKAKELKEQEKKSGKKVHVVLLLDRETGRNSRLLRLERELREEGKSEEEIEEELERERRRLLEEYGVESERERLELIAELFREEGLSDVELIETDREEVEKRIEEIEKQEKEGKPVVLIE | HHEHHHEH | 0.936 | 0.848 | 0.456 |
| N29 | STELAKEYEKKSSEWLEKRKSGSEEEKKLYEKYKKEEQKRKDTKKKSEKYKSTVTITFNTDGTITVKLEEKDEDENISEEEKKEERRTLARETFDAYLQSRKNPNTTVTLTYKVSKKSLEGKDVEGLKDEIIRERRKEIDYYKNELKEPEELINLRKELLYELIDSLYKDDPERRRKLKEELE | HHEEHEHHH | 0.930 | 0.983 | 0.472 |
| N31 | SEELEERLIKEFGSKEEYYSRSVVISESDEESRQLLEEIRQTLQSGLPREEQIERLQELLDKLIEISRNPELLRKYLDSVSKYYGKPLEEVLLEQYEKDSSKSPLEHLYQLFKETADTYYKKEGKKVDVSYEEKDGKKTLIIKE | HEHHHHHEE | 0.930 | 0.983 | 0.472 |
| N15 | SERLERLRSWLEDRRKTASEEERPLLDHSLERLRELEELERKPEEELTEEEKARKEVLRQYSRMLETIESGSSDAPQIREFSKQWSTALDNSLENPGKEQT | HHHHEHEEH | 0.935 | 0.694 | 0.473 |

| | | | | | |
|---|---|---|---|---|---|
| | VTVDSTYSDEFQKLIREYWEALKKLS<br>TESPYKITLKLTTKTKDGSEKTETITIT<br>LSEEQAKDERTLLQSL | | | | |
| N19 | KKEELERLEKEMKESETSESYKKNT<br>VEYLILKKGIKEEDKEKFTEEFNKEVE<br>KLSKEIIEKIKEDRKKGEKSTTIEKTFD<br>LSKSEEELEYESSVYSTAMVKAAEEL<br>GYKNIKELSREMVLAKKVIELAKKKG<br>LKYVKVTTTFKEDGTVEVEIETEDESI<br>LKEAKESLK | HHHEHHEE<br>H | 0.933 | 0.994 | 0.452 |
| N24 | SEEIDKRVEEFWDRVLSENKEYKNW<br>EDVAKAIWDEIEKKAKEEGKTLSEYL<br>YEKLKKVIDENPNKSIEEIKSLFLKELL<br>SSLDSKLSKTMLKELEKTESELESEE<br>EKNEFKNKIYDKLLEEYEKATGEKIEI<br>ELSKDSEGKIKIKVTYTDKNGKEETY<br>E | HHHHHHEE<br>E | 0.938 | 0.880 | 0.456 |
| N17 | DEVWKENIEREKKKYPDWSDEEILIK<br>WTFETLELIIEALEKIGITEEAKELKEE<br>LEELKKKTKNLQDLIDYLEKWLKEEIG<br>EEEYKERSEKEREKSKITDEKLKRW<br>EVTTTLYIDYLKERFEKKHPEYKLNV<br>ETKIEKGKDRKIIVSISNSKTGEKLYET<br>EI | HHHHHHEE<br>E | 0.939 | 0.753 | 0.475 |
| N10 | LDDQIEYYEKLKEEAEKKGYEKLAEK<br>LKKLKETYEEYKKKGNTTENRTERN<br>NKYLEILGTWTDEEIEEYYEVLKEVY<br>KDKSKEYIETHKNHTKYSYDFVKEAK<br>KKGLKVTVTTTIEEKDGKAKLTTEVK<br>GISEELKETAKKAYELYHKLLSEYLKE<br>KKGIKEVETETKF | HHHHHEEH<br>E | 0.935 | 0.869 | 0.467 |
| N13 | DKETYERNKNTINEYLKTKDKSIIESN<br>EETKRVIDILKKEENLKTVEEIIESNMK<br>KYGAKKNSKGELEVTFKTIEEAYEFIR<br>EVSEELSKKFPELKPYLESIEKYSESK<br>EEYINDFVNLLNVAKDYSGRTDYKKL<br>KYSKNSDGSYSITL | HHHEHHHE<br>E | 0.940 | 0.752 | 0.471 |
| N2 | SKKELEEREREKRKELAKSKTEESKK<br>KLLDYYVESELQIEEKKEKGEPESEV<br>EEEKEKLDKEIDEDIDIYYEDKTEEEK<br>KKEKYNFKTEIYREVLRQRALRYASK<br>EELTEEKIRELVRELKKKMDEIRDKY<br>KVKSKEEVDNEVRRMSYETADYLHD<br>EKGLLTEEQRELWRKVVEKEV | HHHHHH | 0.916 | 1.569 | 0.482 |

**Supplementary Table 5. Information about novel 7 helix designs and their MaxTMscore**

| name | pLDDT | ScRMSD | Sequence | Local search MaxTMscore | Foldseek MaxTMscore |
|---|---|---|---|---|---|
| **H7_0** | 0.954 | 0.802 | KKEIEEEKKETEEKVEEDKKSGFLE<br>KSDEEQEKKLTEYLKEKIKKETGIE<br>ELSEETEKEIEESVKRLWKLYKKKY<br>SDREKQLEEMYDSIMKTVLVLKRL<br>KKKYPDLTDEELTESIKLISELEEW<br>DIENPGSSEEEREEQEKKAIEKQYE<br>RYKKKNPDDTEEERKRRKELLEET<br>SKIYREVRKEL | 0.425 | 0.498 |
| **H7_1** | 0.933 | 1.055 | KKETEEELKKWKEEYQKKLEETEE<br>RSESEEELEKTAEELSKEYEEKIEK<br>LEKEKSPLSKEELDKYLKEFKEELL<br>ERLERRRERRREEELEKRRKEYEE<br>TLKKSSDLSEKNKEKLKKRLEKELE<br>YEEKRRELELERKRLEWEKKLLNL<br>PEEEYERRKEEIERKEKEVEEEYK<br>KYIEETTKEDKKEMDDKYEKLKKE<br>GLTEEKRKEYYKYLDDIEKMWEIYA<br>ETTKKEEWKEHYKQIKEHSKKIKEK<br>VEKE | 0.434 | 0.602 |
| **H7_4** | 0.917 | 1.474 | KEKLKKKLNELSEKYKKESDRDEE<br>ELNKSIEIVKRFFFSEEYGEGEKNG<br>SYLLDLRRYMELEAKGTLEEELTEE<br>EKEELKKSGVQLMKDIMRKIIKLRE<br>EYSEEERDIVEDTLRLLTEFIYKTLK<br>EELKEYLDDEEATKKTYETLTKYLK<br>EASEELKKEYSKEEVDEIKKEYEKIL<br>EELKEKEEK | 0.496 | 0.477 |
| **H7_6** | 0.944 | 0.634 | EDEEIDKKLEEYFELVREREERLVK<br>ARQESKTVDEYIKKDREIERERWE<br>KMADLYLKDEEDYEEKKKELSEIHE<br>EELERLEKLDRKNRELVERALEKFT<br>KEEWEEHARRVLERLRERREELKE<br>RLKSLKDSKHKLVYLNSEERYIDYR<br>ETELWYVEKLGVSKEEALELYRERI<br>EIIEESLVRAGVITEEEAEERKKRRI<br>EEMREELE | 0.461 | 0.476 |
| **H7_18** | 0.935 | 1.173 | KKEEEKKKKIEEEVKKLKEKLEKYE<br>KEGKDGRRSLVRDEYEKASGEES<br>EDDITENEVLNKLIKEMSKYEGLNE<br>YEEKKILYGLYSEIEEESKYTSKEEA<br>TLMYVLKLSRLLIERKKNPEETGKEI<br>RKIMERYIEKSEKKIKEKFNNDESIT<br>KEEEEKYLNEELEKQK | 0.426 | 0.449 |
| **H7_26** | 0.915 | 0.956 | SEEEEILQKLEEHLTSFNGKSFSLR<br>QLYEIVKTLEDTYREMIEEYKKTKD<br>LEKSREKAIEEIRKRFQELSEKHSS<br>LDEEELKRIVEKIEKRLKEWTKEKK<br>SIEEVEKYLEELREIRDQLEELLSTL<br>DPELSTEEQIEYWRRELEEAKKRS<br>EEEYKKLLEKYRELVERMEETEEK<br>WEEFL | 0.483 | 0.469 |

| | | | | | |
|---|---|---|---|---|---|
| **H7_27** | 0.916 | 1.760 | REKEKKIEEYTEELRKRLEEEGEED EERIKKRQEETRRILELKYENKELS DEEIYEKYAKEKVKEMIKEGKSLEE VEEELERMEYSASFIEEREKNLSEE EKEEKKSKYEEIKTKALEEYLKEKG KSYEEIYKRKKESLEEQEKRLLEKY TDKEKKEKIKEETEKKLEELDKELE KKKKEEEK | 0.447 | 0.510 |
| **H7_33** | 0.935 | 0.954 | EKEKEEKLNEIFKKSVKELYELEEE KEKLSEEEYEEKYEKILEKTKKEVK EISKEEGESEEKIENDVRLTEMTYK RTRKHIKENYEDIKKNKEKKEEILEE LSKYHYYLFKALRLQKELKYKSEEE KKKKEEEIEKLLKKRNESLEKYLKLL GKTEEEEKNKILESELKVTEDLYKN EKLISKEELEITKQILEKIKEL | 0.456 | 0.475 |
| **H7_50** | 0.944 | 0.703 | EEEETNLLLMKYVIYEEIEETEKRG EEAKKELEKEGKKERLKRTDELVD EKIREDLGIKESKEERDKKKKEAKE YSKEQKKKTEKYLEEHPELKEKVE ELKKKRKELTEELYEEYSRLPKTLE NELKYSDSLISETRRVQIYELEKKG ATLKDLLDTVLKNLELRKKVHREQL RYYRENEAEELGMSREEIEEQERR IEEIIRKIEEYYKETAEEL | 0.488 | 0.464 |
| **H7_55** | 0.912 | 2.088 | ETEDLKKEIKELYEEQIKKVKELTKE VSSLIKTEESREELKKKIEELLKETK ETTEEIIEKIKKAFEKLDKEKKKEEE YKKLIEEILENLSKLNDSYLDLNEKL WEETEEEQREEERKERVKELEKE REKIEKKKEELKKKKKKESKEEKKK LEKKIKKYEERSKVLKENAELLEKD YTSKPEVTDRLKKQELKEKEEEYE EKIEEAKKKGDEEKAKKLEEERKKK KEELEKERKEEVEKTKEEYKEEEK KREERRKEYKEKLEKHKKKHEEKI EELKKKKKVSVEELEKLIKELTEDS EKLTEETLKKVEEEREEYKKK | 0.499 | 0.495 |
| **H7_58** | 0.906 | 1.095 | SEKKEKERIEKEKKRYKESYESGK TTEWEKQTIRILSNKKKLEKSKKTL KEMKEKGELKEEWTEEEVEELIER LEKTIEQLDKYFENKGVKDYESLKK QIEKDYNKIKESEEAKKKNDKSKEE EVKKYEEELKKRVEFEESLSRDKP AEENLLEESKKDLEEAEKYLEDNE EWKEYKKKIEEEEEKKLK | 0.478 | 0.512 |
| **H7_93** | 0.916 | 1.569 | EEKKKKEKEKEELKKKAKSKKEED LKELHEYFVKKKIEIEKLKEKGESEE EIKKKEEEHEKEIDELIEIITEDENEE KKKQKKTETHLKIEEEVHRKRMED RAKKEPITEESIKSDVKDLKKDYEEI RKKYKTKSREEMEKKMRETSEKTA EYLQKEYGILTEEQKKLWRETTEK ES | 0.453 | 0.552 |
| **H7_101** | 0.928 | 1.646 | STEELKKAIEDYSEKLLKGEESDLD EETRKRIEELIKEEIEEYKEYEEYIEK SKDLSEKERVKKVKEKEKEDLLEE WEKESEEERRERRYTERVILETVS EKTGDESLKEDAKIEEEVYEESEEK LKKEKSLSKEEVVEDYIEKLEEKFK | 0.468 | 0.479 |

| | | | | | |
|---|---|---|---|---|---|
| | | | TEKETSYLDHKKKEIEAYKLYEKLRK | | |
| **H7_118** | 0.915 | 1.549 | KTEEKIEKLYEEYLKTELEYVKAEKEGNEKKKLEKYKELLKIEKEIEKLKESGDKKIKESIEKAEKKREEEIEKLPEEERKRIQEIREEIKRERELEKKLEEIKEETKDGSEESKEWKEALETLEEVEKLREERAEKEKKSPKEKEEIYKENLERKEKLIKAKEDPSTRQKYLEELKKELEEEKKKK | 0.449 | 0.463 |
| **H7_122** | 0.943 | 1.016 | IELDVLILVLDTLVKKSKDKEYAEKLWKETKEELKKKKTKEEKIELLKKSIDELLEKEGYSEEKKKDVKYLSEELVKTRKELEKEKTEEEKKEIWEKTEKKLKEYSEYAYELYKEIRKDVLEKTGKIDGYKEYAKEIWEK | 0.463 | 0.453 |
| **H7_137** | 0.955 | 0.622 | SKEEIKKKATEKLEKEVEKISEEYKERWGMDEETLKKIKEETEEDWKRLLELLDKYGTEKYYEYDLETTEKVLTNSYKNRAQREILEKEFGVDKEKFNEKLKKAEEEARKRSEERLEEWKEKKEKSEEELYKEEKERTEKLLEESEELLRRSEELLEKAREGADEKTIEEAEKTLIEAEKLNTEILRDTLYYSALRKARETEKTEEEREEKLEEMEEEIEKTMEEIEKNRQ | 0.474 | 0.575 |
| **H7_141_0_14** | 0.929 | 1.075 | RKKEIEEHAKTWVKYKVLKIKELEGELSEEEKEELKKTEEESDKKSEELEKEKEKYGRKEVRKEWKEAYKKEIEKIEDEKTKAILELVLETEESLRENYYRYKEEKKSEEEILEETKKNFKKIKEKMEKLVKELGVNEETLKKLLREEVEEESEEWKKEGLSEELSEKFKEILDEELEK | 0.491 | 0.562 |

***Supplementary Table 6. Information about selected scaffold for mirror topology.***

| **Fold Name (PDB ID)** | **Fold Type / Function** | **CATH Classification** | **Structural Features** |
|---|---|---|---|
| 1ABE | Response regulator | 3.40.50.2300 | Rossmann-like fold with 3-layer (α/β/α) sandwich topology |
| 4GT8 | Histidine kinase-like ATPase | 3.30.565.10 | Two-layer α/β sandwich, typical of signal transduction ATPases |
| 1QYS | De novo designed fold (Top7) | 3.30.1710.10 | Designed fold with unique α/β arrangement not found in natural folds |
| 2LCI | P-loop NTPase fold | 3.40.50.11230 | Rossmann-related topology with nucleotide-binding elements |
| 2J3W | TRAPP complex component | 3.40.50.620 | TRAPP-domain fold with α/β/α sandwich and curved β-sheet |
| 1SKO | β-lactamase-like protein | 3.30.450.70 | Compact α/β fold found in dynein light chains and β-lactamases |

**Supplementary Table 7. Information about mirror fold designs with experimental validation**

| Rename | sequence | name | pLDDT | RMSD | Max TMscore |
|---|---|---|---|---|---|
| M1 | MYWIVEVEEDENGNIKKVYVKYWTDEELKEGKYSKEELEEWEKREEELIKETKKYEEENKEDESKKELIDDRKRDENGNEKRDIYIPAIKYENKGYVSYLRLTEDTDEEKRKEYLEKSKESLEKILDYLKSKKDKGVTEEDIKKVEKVLEKVKSKI | sketch7_2_T10_17 | 0.849 | 1.600 | 0.445 |
| M2 | KSYRGSIEYDEDGNLNIHKPVLAETEEELKEKLTEEEQKEYEERRKKFSEEYEKAKKLKEQGDKSVYSYHYKLTDENGNTKGEIIYITDEKDKNKLKVYVELYNTENEEDKKKQKEALKKVIEKLTEWYNKYISKNKDEETQKKFEEFLKQLESLL | sketch7_1_T20_56 | 0.877 | 1.630 | 0.444 |
| M3 | MYVYHRLEVDENGKVLYYDSKFYKDEKEYKENISEEEEKEYKDYFEELYNKSKKLKEEKDKNYTIHYNEKEYTTKKDGTKEISYGLVHIEHKDGSLESAIIINPSKNKEEEKKKLKKAKEHLDKVSEYLEKRLDKGVPQSTLNSVKELRDTISKYI | sketch7_0_T20_84 | 0.928 | 0.852 | 0.452 |
| M4 | SKEEKIKEFKEKIEKTKKEIKEKKKDYYLEYEVKDGEAKEPELKEISDEEAEKKIEEYKKYRDELKKSGKEKDYNIKEEEEEEEYNEETKDTSRYVYIHIENKKDKKNRKSLVYYKSNKKTKNEELFKKFEDELEKIYETYKKNL | sketch6_2_T20_50 | 0.899 | 2.131 | 0.392 |
| M5 | KEKKTDEEIRNDLKKAKKNIDKEDISLRLTIDKNGEKKYSYNDESKITEEEKKYYEQTRKKLKEDYEKNKNSDSNYSYYKKERKNSKTGETNSLEILKKKSSDGSSKTEYYLYLKISSSDEDEEFKKEAKEALDESIDEVDNSK | sketch6_2_T10_28 | 0.886 | 2.339 | 0.395 |
| M6 | SLEEKKKEIEELLKRIKEEYESKAFTAYLYYESNGKMEYATTEEELNKEETQKKIKEYKKKAEYYENTEELKKKGVEVEKEEIKRTENGEERTDEYIYIKDKENPENNFTLLTINRKEPITDKEDKEKLKEMIDKYEENLKKQL | sketch6_1_T20_61 | 0.915 | 0.703 | 0.383 |
| M7 | SEQEFLDKAISYLDEARENLTKRPFYYYHSMDEEGNSKSANSEEEYKKKETQDFKERVLKKLEELKNSKDKDKYEIKEEKYEYTEEENGKKKKYKIEYRYARKDGKLKAYYLEIYSSDKMKDEKTIKETKERLDEIKKELESLR | sketch6_0_T20_86 | 0.944 | 1.082 | 0.373 |
| M8 | SLEEVKNAQKELADYLKSDSNSENFSYEEEMTIKNTQGKENKYSRKLEVNKETGEVKTEYNLEWQGSQEQKDNLLQLEKKELEELKKIYEKSSSTDEIITETISMDSSGNVTSDDESVKKSIQDLYNTYKSKLDQGYDINQKEEKLNLSSGKDKSLDYSQSEVSSTNKNLTININIRLPKSAYEHAKNFIINHFQAHKSFLENS | sketch5_2_T20_126 | 0.918 | 0.800 | 0.379 |
| M9 | MIVYLNLLYSTSREGHEEALEMIKETIEELKKGEKNIKKIKLNLEDVDEELKKKTKEKIEESTKKIEEKVEKNNVEKYVVYNIYLSADSSDRESIEEYLKFLEKVTEYILKEIEKGNSVSVTYINFTDSTETEEEKRKKTKEKSEELLKKIEAQSETGIDEVTVLYTPSDEELSNETKKLVEELKEEIRKEREK | sketch4_2_T20_24 | 0.914 | 0.769 | 0.406 |
| M10 | RERRIETDGEVNEEDTKKAIEKLEEANKKLKEGDKYVSYSEISHTSESKEYVEKIRKEWEELNELSKEYYETTGATVSNILYIDNLKKEHKEESEKLTELIKQWRKLLQKAKEKGTKTISFHLVLGDKLESRKKTL | sketch4_1_T20_24 | 0.818 | 1.460 | 0.396 |

| | | | | | |
|---|---|---|---|---|---|
| | DASEKWIKELEEDDEEWDGDELSYYSISKPSKNEELRKKYEEKQEELERLLRESGES | | | | |
| M11 | KNSLEINVNYSYENGKETEIKTEYKLDLPDYDEEKKKEIKEKLEKKLKELKELIEEARKKGIDGLKINLNFKSENVNEEDFDKFYEKLFKKIEETKEKIKEALEKKKPLTLNTNLEYKTEKDKTIYKEEIDEK | sketch 3_2_T20_47 | 0.842 | 1.782 | 0.508 |
| M12 | MFNLKAKLEYDGSKGEEPKLEIEKEIDLNEENKEKLEKETKELEESLKKLKEILEEEKKEGRNDIKVKIEISITNATEDEIDYVIKKLKEITEKIKKLKELLEKRKNGSSEKYEYEEEEEKNNTKIKTKIELK | sketch 3_1_T20_76 | 0.889 | 0.874 | 0.499 |
| M13 | SVKLDIKSDLHSENGKITKDESEYKYEVDEEYKDELGKIVEELKKNVEELKKEVKKLIEEGRKNVKLEFNLKIENTKDEDIEWIKEKLKEINKEVKEITEKLKKLSSNESYKYEKKYEDENIGFKYELKLEVE | sketch 3_0_T20_35 | 0.927 | 0.623 | 0.426 |
| M14 | SEEKEKIEEIKKETKKKIDSGDKNVKTEEQEIELDETTTDEEVVDLVNYIYDSIWENEEYISKKKKKNVSLEYKYKITKTRKDGSKTIIEIEYKSYFKDTYTFSEEEKEKTRERRETMVRTSLETKKDQEEEEEYKSDNYEYKEKLKITHE | sketch 2_2_T20_24 | 0.943 | 0.631 | 0.500 |
| M15 | SELLKETEEAYEKAKKKIEENPEHADITINFNKSGESISLEDIEKSIFKLLDLSSYAVKSKNEDKIKKVNLNINLNVNTTYSDGSTLNVNLNITIENPGDYEEWKKFLDKKKEEIKKTFDEIKETKKPKTYSYSYSDSTTKTSLNLTINYK | sketch 2_2_T10_70 | 0.895 | 1.315 | 0.467 |
| M16 | SESKKLIEEAKKEFDKQLAEGKKEIDIEIEYPKKKYDENYKEALEFIKLNSELRVEYVKKSREAGVEKLTIKTKVKLEPTENSDHKLDYEYEEEMSKKALEKTDEKEYKEKLEKEYKKIEETVESNKDYEYEEEYEDKETGYKYKKKVEIK | sketch 2_0_T20_65 | 0.933 | 0.755 | 0.469 |
| M17 | SKEMEEFNKEIEEYKEELKKSKEEINIKFPYKKYTENVSYEELKTEKTKLIDLLKEIYNTTKEKKLKKVKFEYKREYSYKTEDGSEVYIKLEISASNLNKETNYEEKTNDFEKKFKSSLDKAEKSTEDLSEEYEYKDDKSGLKLNYKLERK | sketch 2_0_T10_100 | 0.901 | 1.111 | 0.428 |
| M18 | KRTHSLNLDIDIDFKGKDKNINKEEQNEIKETLEEQKNYLKTLLENADTGKTVSYNSYKKEYKSGLTINKSGDDISSDSTETKTLSLNINYDDKKTGNKLNINLSLKLTSIKDSRETIEKALENVEKIEKWIEESLSTGKSYKYSYSYTEEK | sketch 1_2_T10_81 | 0.908 | 1.674 | 0.412 |
| M19 | SNKKTIKSEVDITRKKTDPKRDKEMKEKTQQKLDQNESNTNNYLNNIDNYDKESETENKTEETEDFTINNNTTYTKEKDNKKYTYNQKTTITDKKSGSKVNINITITNTDKTWSDEFKQKYNNSYQNIKNSINTSFNTGKNIKKNNEESLEY | sketch 1_0_T20_117 | 0.881 | 1.977 | 0.502 |
| M20 | MKYYESIIKIGVSEEEQKELLEKTKERIEELRKKIEEGSDEEYSYSEISYDESLSEEHKKILERIKETNKKIKEKLKEGDVYLSIYITSSTELVKKEDLEEILEKNEEYLKKLEEGEEKPYYSYTKLKGVSKELEESAKELEETVREVYNKT | sketch 0_2_T20_89 | 0.844 | 0.875 | 0.450 |
| M21 | KIRSTLLIILKASEETLKETIEIIEEELERLKELREQGRRERTSTVKLTLSDDVPEEDREWLEETIREIKEELDKLSDDDIPGPIIVTYSNDGVSEEEKEEGKRTLEEYREELRERKENIPSRILLSNLESEELEKELREKYEELDQLLRSL | sketch 0_0_T20_43 | 0.874 | 0.735 | 0.430 |
